# EMPIRE: The Ellipse Model for Phylogenetic Inference of Range Evolution

**DOI:** 10.64898/2026.04.23.720387

**Authors:** Sarah Swiston, Sean McHugh, Michael Landis

## Abstract

Many phylogenetic models of historical biogeography exist for describing how lineages move and evolve over time. Here, we present the Ellipse Model for Phylogenetic Inference of Range Evolution (empire), which models the movement and splitting of species range ellipses in continuous space, summarizing important attributes of each range, such as its position, size, and orientation. The framework allows us to reconstruct ancestral range ellipses, investigate rates governing important processes like movement, expansion, and elongation, and examine the spatial context of speciation, including asymmetric range inheritance at cladogenesis. We apply empire to the Australian Sphenomorphinae, a group of skinks whose diversification has coincided with substantial climatic change over the past ~36 million years. We find that speciation events are positively associated with aridification, while daughter lineages post-speciation do not tend to show evidence of ecological partitioning.

---

Evolutionary biologists are interested in how species move and evolve in space. Knowing how the geographical ranges of species, and the species themselves, have spread and split over time would help us understand how lineages tend to interact with the environment and predict how lineages will respond to future climate change. In order to investigate the spatial history of clades of closely related species, researchers have developed numerous models of historical biogeography that use information about the phylogenetic relationships and spatial distributions of present-day species to reconstruct ancestral locations and infer rates of important processes, including dispersal, extinction, and speciation.

However, realistically modeling the phylogenetic evolution of species ranges is challenging. One reason for this is that actual species ranges are complex, and experts have different ways of representing them at different scales, recognizing that different representations can be more or less useful for different purposes. For example, biologists often represent expert range maps as polygons, even though those polygonal range maps may fail to exclude all areas where a species is absent (e.g., by including interior lakes where land-dwelling lizards could not live). Nonetheless, these maps remain useful for understanding where a species can be found, or assessing how many species may be present in a particular location, at broader regional scales. In the context of phylogenetic models, we also need to represent ranges in a manner facilitating our description for how species ranges evolve over a tree, both anagenetically (between speciation events) and cladogenetically (during speciation events). In general, high-resolution representations (e.g. polygons, sets of occurrences) can be more difficult to work with mathematically than low-dimensional representations (e.g. regions). This has led to the development of two main categories of phylogenetic models of biogeography: discrete-space models and continuous-space models. Each category has its own benefits and drawbacks.

Discrete-space models, the first category, subdivide the space in which a lineage is evolving into several regions, where each species may be present or absent in any particular region. Models of this kind include DIVA (Ronquist 1997), DEC (Ree et al. 2005; Ree and Smith 2008), DEC+j (Matzke 2014), BIB (Sanmartín et al. 2008), DAISIE (Valente et al. 2015), GeoSSE (Goldberg et al. 2011), and FIG (Landis et al. 2022; Swiston and Landis 2025). Although each model has different properties, they enable biogeographers to estimate dispersal rates (and sometimes extinction and speciation rates) for species distributed among the involved regions. Many of these methods can also be used to reconstruct which regions any ancestor most probably occupied. Such models are most useful in systems where the regions represent distinct entities of interest, such as islands, entire continents, or habitat patches. However, in many systems, a useful regionalization does not immediately present itself, and the consequences of splitting space into discrete regions are poorly understood. For example, species ranges rarely occupy entire continents in continental biogeography, yet a species occupying even a small portion of a continent must still be labeled as ‘present’ or ‘absent’ for that region. An attractive solution would be to reduce the sizes of regions to have a finer-scale resolution, but unfortunately, increasing the numbers of regions increases the number of possible ranges (the state space) and, often, the number of model parameters (Ree and Sanmartín 2009).

Continuous-space models of range evolution, the second category, use a point (e.g. the centroid of geospatial occurrences) or a set of points (e.g. of all geospatial occurrences) to represent a species range, and describe how it evolves among different species over time. Models of this category generally use a two-dimensional diffusion process, representing how the range of a taxon (represented as a point) changes its latitude and longitude over time. Examples include standard random walks (Lemmon and Lemmon 2008), rate-heterogeneous random walks (Lemey et al. 2010), random walks that account for sampling uncertainty from the species range (Quintero et al. 2015), random walks on spheres (Bouckaert 2016), and random walks under changing paleogeographic conditions (Arias 2024). This class of models shows great promise, since species ranges exist in continuous (not discrete) space. That said, continuous-space models, such as those mentioned above, do not track important properties of species ranges as they evolve over time, such as range size or shape, which are captured by discrete-space models (albeit in very low resolution). In addition, and also unlike discrete-space models, current continuous-space models assume that ranges are inherited equally by both daughter lineages after a speciation event, limiting their ability to make inferences about the spatial context of cladogenesis.

A third class of “hybrid” models instead use fine-scale grids to approximate continuous space, describing ranges as a collections of occupied grid cells. Models within this class are generally simulation-based, such as DREaD (Skeels and Cardillo 2019) and gen3sis (Hagen et al. 2021). As is the case with most discrete-space models, parameters for these models usually control the behavior of species occurrences moving and diversifying among grid cells, rather than describe the properties of whole ranges. Translating between grid-based estimates and emergent properties of evolving species ranges (size, orientation) is not trivial, and defining mathematical rules for how cladogenetic events occur in continuous space is challenging. SEAMLESS (Albert et al. 2017) is another exemplary, but simulation-based, model that uses the stochastic appearance and erasure of barriers over time to generate interpretable phylogenetic patterns of species ranges. Any such simulation-based models, however, do not have have known likelihood functions, and rely upon likelihood-free estimation methods (e.g. Approximate Bayesian Computation or machine learning) to analyze real data. In addition, such models do not provide methods for performing ancestral state reconstructions, which generally requires likelihood-based inference.

To our knowledge, no likelihood-based models describing the evolution of species ranges – with locations, sizes, and orientations, and that can be inherited asymmetrically during cladogenesis – in continuous-space currently exist. Here, we present the Ellipse Model for Phylogenetic Inference of Range Evolution (empire). It is an ellipse-based continuous-space model that lets us describe three evolving range properties which are interesting to biologists and essential for reconstructing spatially-explicit cladogenetic events at nodes in the phylogeny: location, size, and orientation. empire uses ellipses to summarize species ranges, which evolve like lineage traits over time. While ellipses do not capture the exact boundaries of species ranges, ellipses offer a mathematically tractable representation of the extent and orientation of ranges. Using empire, we can use data about the distribution of present-day species and their inferred relationships to investigate how lineages move and split over time. We can estimate important model parameters describing how lineages change location, shape, and size, both anagenetically and cladogenetically. empire can also be used reconstruct ancestral ellipses at the nodes in a phylogenetic tree. To infer model parameters and ancestral ranges, we developed a new data augmented Bayesian Markov chain Monte Carlo method to fit empire to data.

We assess the usefulness of EMPIRE through the analysis of simulated and empirical data. We first measure the accuracy and precision of EMPIRE using data simulated under different scenarios to validate conditions where the model and method performed reliably. After validation, we used EMPIRE to investigate the historical biogeography of Australian Sphenomorphinae, a lizard clade containing nearly 280 recognized species (Singhal et al. 2025; Torkkola et al. 2026). The phylogeny used in our analyses contains 218 of these species (Title et al. 2024). This study system is ideal for the application of EMPIRE, as it contains enough species to inform rate estimates, but not so many species that the data augmentation procedure used to calculate likelihoods becomes intractable. Additionally, the model assumes that geography is even and traversable, so a continental system with few barriers is preferable. These skinks have diversified and spread across Australia over the course of ~36 MY, experiencing the continent’s substantial increase in aridity over that time. It has been hypothesized that this environmental change impacted the evolution and movement of the skinks (Rabosky et al. 2007). By reconstructing the ancestral areas at nodes in the phylogeny, we investigate whether speciation events are spatially coincident with areas of aridification, and whether daughter lineages demonstrate ecological partitioning. We also examine the rates at which ranges move through space, how they change in orientation, and how they tend to grow between speciation events.

## Materials and Methods

### Range Ellipses

The EMPIRE model represents species ranges with ellipses that move through space, change in shape and size, and split at cladogenesis (Fig. 1a). Because the ellipses in the EMPIRE model evolve stochastically over time, they are defined in a somewhat unconventional manner. Ellipses can be mathematically described in terms of major and minor axes, and an angle of rotation. However, due to the nature of rotation, there are an infinite number of ways to describe any ellipse using these attributes. Constraining the amount of rotation does not solve the problem; regardless of where the constraints are placed, similar ellipses will be treated as being very different (Supplementary Fig. 6).

**Figure 1.**
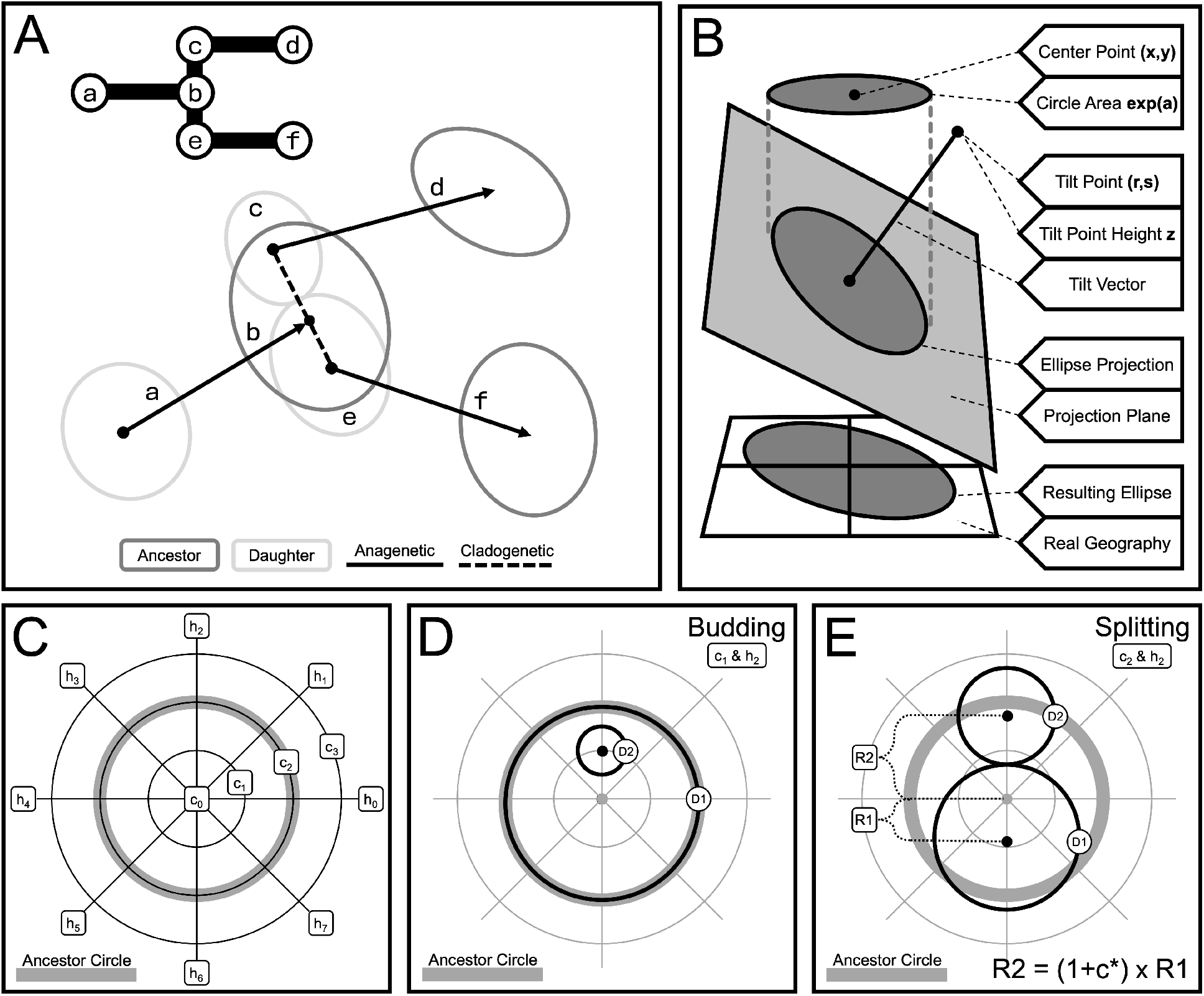
Description of the EMPIRE model. **A)** An example of a range ellipse evolving through time, including anagenetic movement (solid arrows) and a cladogenetic event (dotted lines). Ancestral ellipses at phylogenetic nodes are shown in dark gray, while new daughters are shown in light gray. A phylogeny is also provided. **B)** A visual representation of a circle projected to create an ellipse. The circle, with centroid coordinate (*x, y*) and area *exp*(*a*), ‘casts a shadow’ on the tilted plane defined by the tilt vector (*r, s, z*), which is ‘flattened’ onto the plane representing real geographic space (latitude and longitude). **C)** The grid system for cladogenetic scenarios, including concentric circles *c* and direction lines *h*. **D)** The budding cladogenetic scenario. One daughter (*D*1) inherits the whole ancestral range, while the other daughter (*D*2) inherits a small range, located at the intersection of concentric circle *c* and direction line *h*. **E)** The splitting cladogenetic scenario, in which daughters partition the ancestral area. Partitioning may be symmetrical or asymmetrical, with the degree of asymmetry governed by the concentric circle *c*. Centroids relocate along the axis *h* such that the daughters are adjacent and non-overlapping.

Instead, EMPIRE uses projection to describe ellipses (Fig. 1b). A circle, when projected onto a tilted plane, will create an ellipse. This solves the rotation problem, as there is only one circle and two such tilted planes that create any given ellipse. Any tilted plane can be defined by a perpendicular vector in three dimensions. While any point along this vector could be used to define the plane, fixing one of the dimensions to an arbitrary value (in this case, vertical height, *z*) allows the plane to be uniquely described by a single point. Therefore, any ellipse can be uniquely defined by a projecting circle with a particular location and size, and a set of only two tilt points with symmetrical coordinates. During analysis, both tilt points defining the same ellipse can be examined without much difficulty.

We use the variables *x* and *y* to describe the coordinates of the center point of a range ellipse, and use the variables *r* and *s* to describe the coordinates of the tilt point which controls ellipse oblongness, where *r* and *s* use the same coordinate system as *x* and *y*. We use *a* to denote the log area of the projecting circle which controls ellipse size. A full list of model characters and parameters describing ellipses and their evolution can be found in Supplementary Table 1.

In reality, species ranges are not elliptical, so range data about extant species must be converted into a set of ellipses prior to analysis. There are several possible methods for transforming a set of occurrence points or an expert range map into an ellipse. For example, a search algorithm might identify the smallest possible ellipse that contains all occurrence points, or that circumscribes the entire range polygon. An alternative method used in this paper is to create confidence ellipses around a set of points. These points could be occurrence points, points drawn uniformly from the interior of a range polygon, or points sampled from an expert range map with occurrence probabilities. This method has the benefit of removing outliers, so ranges are not needlessly expanded to include large areas which are mostly unoccupied. Additionally, confidence ellipses can be easily computed using the covariance matrix of the set of points; the directions of the major and minor axes are determined by the eigenvectors of the covariance matrix, and their lengths are determined by the eigenvalues and the desired degree of confidence.

### Range Evolution

Modeling the stochastic evolution of the circles and tilt points used to create ellipses is relatively straightforward. Each ellipse is defined using a set of five evolving characters: {*x, y, r, s, a*}. Ranges evolve along branches in-between speciation events (anagenesis) and at the ends of branches during speciation (cladogenesis). After modeling the ellipses and the processes that govern their anagenetic evolution, we introduce events to describe range splitting at cladogenesis (Fig. 1c–e. We assume that the true dated phylogeny is known, and only observed cladogenetic events are modeled, ignoring the possibility of hidden speciation and extinction events. (We later show results from a sensitivity analysis to assess the influence of such missing taxa). In addition, we assume ranges evolve in a flat and unbounded landscape so that the likelihood function is tractable. The statistical methods used to compute model likelihoods while accounting for the unequal inheritance of ranges is similar to the Asymmetric Brownian Motion model described by Gaboriau et al. (2025). We will define our events in ‘circle space’ (happening to the circle that projects to form the ancestral ellipse), and then project those events into real space (the actual plane where elliptical species ranges are evolving) using functions *f* (*r, s, a*, …) (Supplementary Fig. 2 and Supplementary Table 2).

#### Anagenetic changes

We use characters *x* and *y* to model the latitude and longitude of the center point of the species range ellipse. These coordinates evolve under univariate Brownian motion (Felsenstein 1985) with volatility *σ*_*x*_ and *σ*_*y*_, respectively (i.e., the square-root of the diffusion rate, *σ*^2^). The log area of the projecting circle is represented using the character *a*, which evolves under a univariate Ornstein-Uhlenbeck (Hansen and Martins 1996) process with volatility *σ*_*a*_, long-term mean *µ*, and rate of mean reversion *κ*. The Ornstein-Uhlenbeck process is used in place of Brownian motion for size because we expect many daughter ranges to be reduced after cladogenesis, requiring a trend toward expansion to explain the general constancy of range sizes over large spans of time.

Changes to the tilt point coordinates used to define the elongation of the ellipse, *r* and *s*, are also modeled with Brownian motion. These characters evolve under univariate Brownian motion with volatility (i.e., the square root of the diffusion rate) *σ*_*r*_ and *σ*_*s*_, respectively. The height of the tilt point, *z*, is fixed to an arbitrary value (we use the value 10 in our analyses). The same *z* value must be used to generate the data ellipses for extant species and to analyze that data. Changing this height will impact the relationship between *r, s* motion and ellipse oblongness. Essentially, it defines the *scale* for *r* and *s*. For instance, a very large value of *z* requires similarly large values of *r* and *s* to generate oblong ellipses, and rates of evolution for *r* and *s* will be greater. Alternatively, a very small value of *z* requires similarly small values of *r* and *s* to avoid generating very oblong ellipses, and rates of evolution for *r* and *s* will be smaller. These values can only be interpreted in the context of one another.

#### Cladogenetic events

EMPIRE models cladogenetic range evolution for divergence events represented in the phylogeny. We assign a base probability for each cladogenetic event type over all nodes, and then reconstruct the posterior distribution of events at each internal node. At each cladogenetic event, either the left or right daughter is selected to be the first daughter lineage, *D*1, with equal probability; the other daughter is assigned *D*2. We call the configuration of the daughters at a node *d* ∈ *{*0, 1*}*. A value of *d* = 0 indicates the left daughter is *D*1 and a value of *d* = 1 indicates the right daughter is *D*1. Each daughter type (*D*1 or *D*2) behaves differently during cladogenesis, so estimating the daughter configuration is relevant. The event is also assigned indices indicating a budding or splitting mode *m* ∈ *{*0, 1*}*, a concentric circle *c* ∈ *{*0, 1, 2, 3*}*, and a direction line *h* ∈ *{*0, 1, 2, 3, 4, 5, 6, 7*}*, assuming we have four concentric circles and eight direction lines (Fig. 1c–e). A value of *m* = 0 indicates a budding event and a value of *m* = 1 indicates a splitting event. The intersections of the concentric circles and direction lines represent a grid of pre-defined, discrete spatial scenarios. Each concentric circle *c* and each direction line *h* is associated with an actual predetermined value *c*\* or *h*\*, describing the radius of the concentric circle relative to an ancestral circle at cladogenesis and the angle in radians of the movement of daughter *D*2, respectively. Any number of concentric circles and direction lines could be defined, but we would caution that a finer grid is not always better; it increases the run time of the analysis, and there may not be enough signal in the dataset to distinguish between many similar scenarios. For our analyses, we use four concentric circles (relative radii) 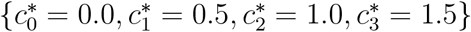 and eight direction lines 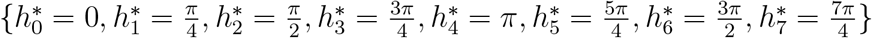.

We represent the base probabilities of these events using *ρ*_*m*_, *ρ*_*c*_, and *ρ*_*h*_. In our simple model, the base probabilities of all modes, concentric circles, and direction lines are equal and independent of one another. In a more complex model, a prior (e.g. a Dirichlet distribution) could be assigned to the base event probabilities. We can call the overall cladogenetic scenario at a node **w**_*node*_ = *{d, m, c, h}*, and the set of all cladogenetic scenarios on the tree **W**.

In the budding scenario, the daughter assigned *D*1 inherits the entire ancestral range. Therefore, the *{x, y, r, s, a}* values are all preserved for *D*1. The daughter assigned *D*2 inherits a new range. The new range is given a small, fixed value for *a*, set equal to *α*. The *α* parameter could be estimated, but in our analyses, we fix *α* to a value of −5, which corresponds to a radius of ~ 5 kilometers. This value represents an arbitrarily small range for a nascent species, and is only meaningful in context of the OU process governing the evolution of the log area character *a* (the log area of the projecting circle). The new range is also given a new center location. Its location relative to the ancestral circle is defined by the intersection between the concentric circle *c* and direction line *h*. Its total position in circle space also depends on the size of the ancestral circle, *a*. Note that *c, h*, and *a* relate to the circle which is projected to form the ancestral ellipse, so the actual *x* and *y* values for *D*2 are determined via projection functions *f* (*r, s, a, c*\*, *h*\*). These functions also depend on the oblongness of the ancestral ellipse, as defined by *r* and *s*. The new ellipse retains its oblong properties, so *r* and *s* values are preserved after cladogenesis. The maintenance of oblongness after speciation is based on the assumption that the forces which cause oblongness in an ancestor should still act upon the descendant species, although alternative models could be constructed.

In the splitting scenario, neither daughter inherits the entire ancestral range. Instead, they partition the ancestral area *a*, with a degree of asymmetry based on the selected concentric circle *c*. Keep in mind that while the total covered area remains the same between ancestor and daughters, the daughter ellipses will not cover precisely the same geographic space as the ancestor. The area of daughter *D*1 is larger than the area of daughter *D*2 by a factor of (1 + *c*\*)^2^, where *c*\* is the relative radius of the concentric circle *c*. The center point of daughter *D*2 relocates in the direction associated with the direction line *h*, and the center point of daughter *D*1 relocates in the opposite direction, such that the daughters are non-overlapping but adjacent. The magnitude of relocation may also be asymmetrical, with the movement of *D*2 being greater than the movement of *D*1 by a factor of (1 + *c*\*). Under this model, the splitting scenario with maximum asymmetry is most similar to a peripatric budding scenario. Again, note that *c, h*, and *a* are related to the circle which is projected to form the ellipse, so the actual *x* and *y* values for *D*1 and *D*2 are determined via projection functions *f* (*r, s, a, c*\*, *h*\*). These functions also depend on the oblongness of the ancestral ellipse, as defined by *r* and *s*. Both daughters retain their oblong properties, so *r* and *s* values are preserved.

### Inference Procedure

Analyses under EMPIRE are performed in a Bayesian framework using Markov chain Monte Carlo Metropolis et al. (1953); Hastings (1970). The procedure relies on data augmentation, assigning values to internal nodes of the tree using MCMC proposals, then conditioning on those values for total likelihood calculations (Robinson et al. 2003; Landis et al. 2013).

#### Data augmentation: discrete events

The first step is to assign cladogenetic scenarios **w**_*node*_ = *{d, r, s, a}* to internal nodes and calculate the probability of this history according to base event probabilities *ρ*_*m*_, *ρ*_*c*_, and *ρ*_*h*_. Under a simple model with uniform probabilities for events, updating the cladogenetic scenarios at nodes does not change the probability of the cladogenetic history. While the probability of a cladogenetic scenario occurring at a node is not based on other model features (like stochastic process rates or root states), we will condition on Θ to simplify the notation, where Θ incorporates all model parameters with fixed values or priors, as well as the time-calibrated phylogenetic tree. Fixed parameters include the tilt point height *z* = 10, the small range size *α* = −5, and the uniform base event probabilities (*ρ*_*m*_, *ρ*_*c*_, *ρ*_*h*_). Estimated parameters with assigned priors include starting states (*root*_*x*_, *root*_*y*_, *root*_*r*_, *root*_*s*_, *root*_*a*_) and parameters associated with stochastic processes (*σ*_*x*_, *σ*_*y*_, *σ*_*r*_, *σ*_*s*_, *σ*_*a*_, *µ, κ*). We calculate the total probability of the data-augmented history of cladogenetic events **W** by taking the product of the per-node probabilities:

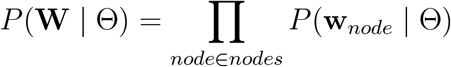

#### Data augmentation: continuous characters

Next, we assign values of *r, s*, and *a* to ancestors at nodes and calculate the probability of this history given the cladogenetic scenarios. These values at a given node, which we will call **v**_*node*_ = *{r, s, a}*, represent the values of continuous ellipse characters *going into* the cladogenetic event (the ancestral values). Values *coming out of* cladogenetic events (the daughter values) are deterministically calculated from the ancestral values **v**_*node*_ and the cladogenetic scenario **w**_*node*_ (see Fig. 2 and Supplementary Table 2). Under the current model, *r* and *s* do not change across phylogenetic nodes under any scenario, while *a* has the potential to remain the same (under the budding scenario for *D*1, decrease (under the splitting scenario), or be fixed to a small value *α* (under the budding scenario for daughter *D*2).

**Figure 2.**
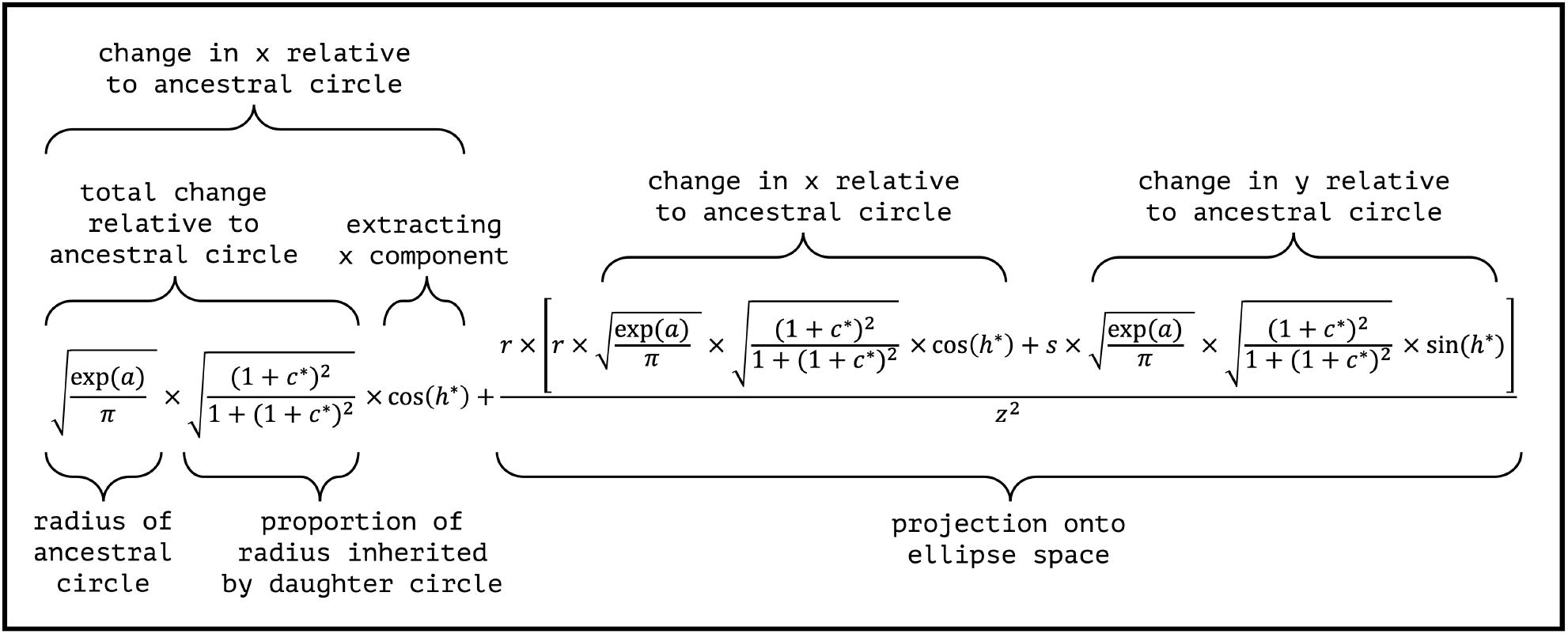
An example projection equation. This equation calculates the change in *x* for daughter *D*2 after a splitting cladogenetic event, given data-augmented values *{r, s, a, c*\*, *h*\**}*. Note that the equation is based on simple trigonometry and does not involve costly operations, but does require exponentiation of *a*, which may cause overflow issues if scale is not properly accounted for. This is the only part of the procedure that takes place outside of log space. (Eq. *f*_3_ in the supplementary material.)

The probability for a single branch can then be obtained, since *r* and *s* evolve under Brownian motion, and *a* evolves under an Ornstein-Uhlenbeck (OU) process. We index each branch by its descendant node, referring to the parent of that node using *pa*(*node*). The starting state for each branch is determined by the values at the parental node **v**_*pa*(*node*)_ (possibly transformed according to the discrete scenario at the parental node **w**_*pa*(*node*)_), and the ending state for the branch is **v**_*node*_). Finally, we know the branch length *l*_*node*_. These values enable us to calculate branch-wise transition probabilities, using Brownian motion for *r* and *s* (Felsenstein 1985) and an OU process for *a* (Martins 1994).

As *r* and *s* are treated identically, we will provide the equations for *r* only. For a Brownian process with start state *r*_0_, end state *r*, rate 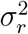, and branch length *l*, the probability is as follows:

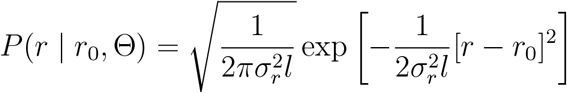

For an OU process with start state *a*_0_, end state *a*, rate 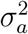, long-term mean *µ*, rate of mean reversion *κ*, and branch length *l*, the probability is as follows:

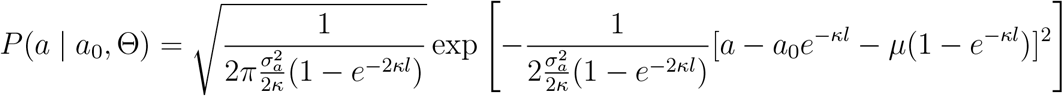

When using real data, we also have to select *r* and *s* for the tips, as two different *r, s* pairs can describe the same ellipse. We calculate the total probability of the data-augmented history of continuous values at the nodes, **V**, by taking the product of the per-branch probabilities:

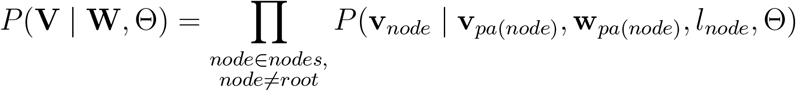

Then we can calculate the joint probability for the data-augmented histories, **W** and **V**. This is simply the probability of the cladogenetic history **W** multiplied by the probability of the data-augmented continuous values **V** given that history.

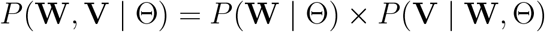

#### Centroid movement

Next, we calculate the probability of observed centroid location values *x* and *y* given the cladogenetic scenarios and values of *r, s*, and *a* at nodes. Since *x* and *y* are treated identically and independently, we will discuss the calculations for *x*. We use a standard phylogenetic variance-covariance matrix, as the variance of *x* does not change across phylogenetic nodes. However, because the value itself does change at cladogenesis, we need to update expected values for *x* at the tips of the phylogeny by traversing each node and implementing one of several transformation functions for each daughter’s descendant tips (see Fig. 2 and Supplementary Table 2). This is a pre-order tree traversal beginning at the root of the phylogeny, where each node must be visited before any of its descendants can be visited and all nodes must be visited once. Expected values for all tips are initially set equal to the root value *root*_*x*_. As nodes are traversed, transformations are applied to all descendant tips; one transformation is applied to all descendant tips of the left daughter, and a different transformation is applied to all descendant tips of the right daughter. The traversal is complete when all internal nodes have been visited. The procedure adjusts the expected values of *x* (and of *y*, analogously) for all species to account for cladogenetic range inheritance.

These transformations depend on the ancestral values of continuous characters **v**_*node*_ = *{r, s, a}* and the cladogenetic scenario **w**_*node*_ = *{d, m, c*\*, *h*\**}*. In this case, *c*\* and *h*\* are quantities associated with particular selections of *c* and *h*. Specifically, *c*\* is the relative radius of the concentric circle when compared to the original circle, and *h*\* is the angle of the direction line in radians. Note that *c* and *h* are related to the original circle that is projected to form the ellipse. Therefore, the *x, y* locations of the daughter ellipses will be subject to projection, and *h*\* cannot be interpreted as an absolute direction of cladogenesis.

After calculating the expected values of *x* and *y* at the tips of the phylogeny, given the data augmented history, we calculate the probability of observed values **X** and **Y**. Probabilities for **X** and **Y** are calculated identically, so we will provide the equation for character *x*. Let **X** be the vector of observed values, **E**[**X**] be the vector of expected values, 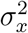 be the diffusion rate of Brownian motion along the x axis, **C** be the phylogenetic variance-covariance matrix defined by the tree structure, and *n* be the number of tips in the phylogeny. We calculate the likelihood of observing the data using the following equation (Harmon 2018):

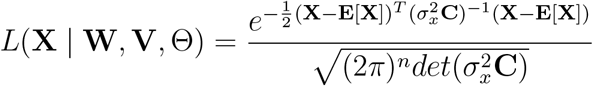

Because characters *x* and *y* evolve independently, the joint probability of the observed values **X** and **Y** equals the product of their marginal probabilities.

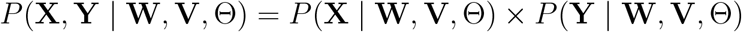

Finally, we calculate total model probability. This is the joint probability of the data-augmented histories **W** and **V** multiplied by the probability of the observed values **X** and **Y** given that history.

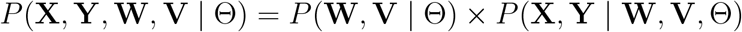

#### Markov chain Monte Carlo

Using this procedure to calculate likelihoods, we fit the model using Bayesian inference by performing MCMC proposals on each of the model components Θ, **W**, and **V**. All non-fixed parameters and data-augmented continuous characters are examined using sliding proposals. The hyperparameters for these sliding proposals are tuned during burn-in, targeting an acceptance ratio between 0.2 and 0.4. For discrete cladogenetic scenarios, symmetrical MCMC moves propose the alternative daughter configuration, the alternative cladogenetic mode, an adjacent direction line *h*, or an adjacent concentric circle *c* (the maximum and minimum *c* are also considered adjacent to one another). Moves are also performed on tip values of *{r, s}*, proposing inverse values *{*−*r*, −*s}* which describe an identical range ellipse. During each MCMC iteration, a single proposal is chosen according to a set of proposal weights, which can be modified by the user. If a proposal on a data-augmented value from **W** or **V** is selected, a target node is then identified, and individual proposals are made on that node along with any extant parent, child, grandchild, and great-grandchild nodes in a randomized order.

This data-augmentation procedure naturally results in ancestral state reconstruction for ellipse characters *r, s*, and *a* at internal nodes in the phylogeny. To reconstruct centroid coordinate *x* or *y* at the nodes, we can draw from a multivariate normal distribution across all nodes, conditioned on the observations at the tips and estimated characters at the root, accounting for the impact of cladogenetic events with a tree-traversal procedure like the one used to calculate model probabilities. This is done after each saved iteration, creating reconstructions for these characters that are consistent with each explored history of cladogenetic events.

### Software

The EMPIRE model is implemented in an R package. The EMPIRE R package is available on GitHub: https://github.com/sswiston/empire. It can be installed using the REMOTES package (Csárdi et al. 2024) by entering: `remotes::install_github(“sswiston/empire”)`.

The EMPIRE package uses four custom classes to manage data and perform analyses. The first custom class is the `detailedTree`, which is designed to hold a phylogenetic tree of class `phylo` from the ape package alongside other important information about the tree structure (Paradis and Schliep 2019). The second custom class is the `dataTree`, which contains one `detailedTree` object, ellipse tip data and internal node data, and a `ellipseParam` object (the third custom class) with values for each model parameter. The fourth custom class is the `ellipseMCMC`. This class contains one `dataTree`, accepts input related to model priors and analysis conditions, and maintains information about model probabilities throughout the MCMC analysis. The `run_MCMC()` function can be used to run the `ellipseMCMC` object, and includes arguments for the duration of the MCMC analysis, the number of burn-in iterations, and thinning.

An analysis using the EMPIRE package results in a series of log files in .*tsv* format. These log files contain information about all model parameters (*model_log*.*tsv*), cladogenetic scenarios at internal nodes (e.g. *d_log*.*tsv*), and continuous values at internal nodes (e.g. *r_log*.*tsv*). Using this simple format allows the MCMC output to be easily read into R, as well as other programs. The EMPIRE package provides several post-processing and plotting functions. For all simulation and empirical analyses in this study, post-processing was performed in R.

### Simulations

To investigate the performance of the EMPIRE model, we performed several sets of simulations. The first set of simulations was used for a coverage analysis. Data was simulated and analyzed under EMPIRE using an identical set of priors. If the simulation and inference machinery is working properly, we would expect that the 95% highest posterior density (HPD) interval for any estimated character or parameter should include (‘cover’) the true simulating value 95% of the time. For our coverage analysis, we simulated 200 datasets using EMPIRE. First, phylogenetic trees were generated using the R package phytools (Revell 2024). These trees were constrained to have a height of 40 time units (similar to the Australian Sphenomorphinae, with a crown age of ~36 Ma) and 20–250 taxa. Because EMPIRE does not model the branching process, the trees were simulated under a pure-birth model, and trees that did not contain an appropriate number of taxa were simply rejected. For each simulated tree, we then drew root states and model parameters from a common set of priors. Values for the volatility (i.e., the square root of the diffusion rate) of centroid and tilt point evolution (*σ*_*x*_, *σ*_*y*_, *σ*_*r*_, *σ*_*s*_) were drawn from a uniform distribution between 0 and 1. Root node coordinates (*root*_*x*_, *root*_*y*_) were drawn from a normal distribution with mean 0 and standard deviation 1. Root elongation coordinates (*root*_*r*_, *root*_*s*_) were drawn from a normal distribution with mean 0 and standard deviation 5. Root log area (of the projecting circle) *root*_*a*_ was drawn from a uniform distribution from −5 to 10. For the OU process governing the evolution of log area (of the projecting circle), the volatility parameter *σ*_*a*_ was drawn from a uniform distribution between 0 and 5, the long-term mean *µ* was drawn from a uniform distribution between −5 and 10, and the rate of mean reversion *κ* was drawn from a uniform distribution from 0 to 5. Fixed parameters *z* and *α* were set to 10 and −5, respectively. After drawing our root states and model parameters, we assigned discrete cladogenetic scenarios to nodes, assuming that daughter configuration, cladogenetic mode, concentric circle, and direction line were independent and all options were equally probable. Then we simulated the evolution of ellipse characters from the root of the phylogeny to the tips. These simulated datasets were then analyzed using the same priors to assess coverage. Each MCMC was run for 2,000,000 generations, with 100,000 generations used for burn-in to tune proposal widths. Typical run times were 1–2 weeks. After analysis, estimated root states and parameter values were compared to the true values used in simulation, and coverage was calculated for each root state and model parameter using 95% HPD intervals (Fig. 3).

**Figure 3.**
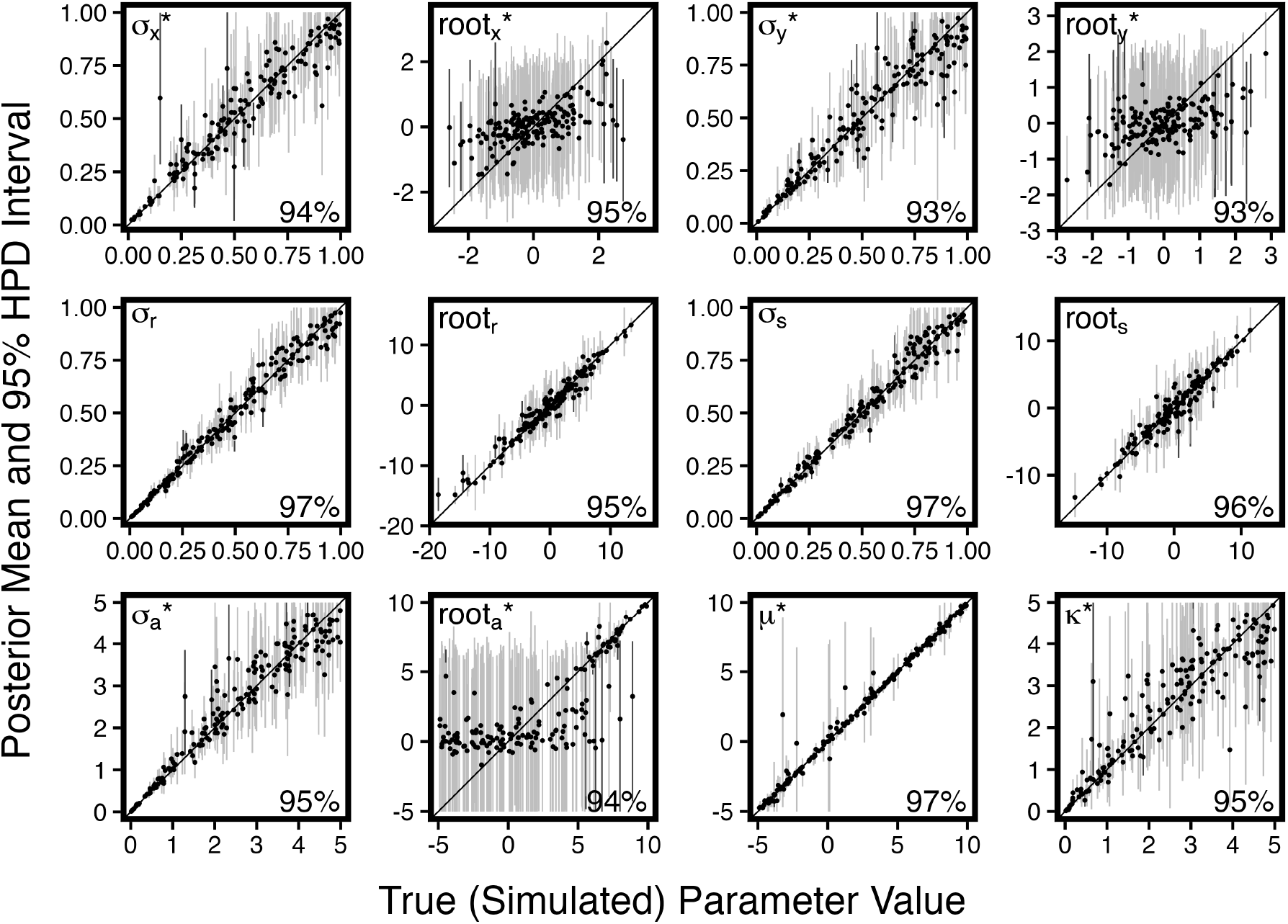
The results of a coverage analysis for each root state and model parameter describing the anagenetic evolution of ellipses. Coverage values (lower right) are calculated using 95% HPD intervals, and should therefore target 95%. Intervals which cover the truth are shown in light gray, while intervals that do not cover the truth are shown in dark gray.

We also performed sensitivity analyses to show how EMPIRE performs under non-ideal conditions. First, we assessed the impact of improperly-defined modern-day ellipses. Real species ranges are often difficult to estimate, particularly if our knowledge of a species is limited. Ellipses also imperfectly summarize the properties of species ranges, and some ellipses are better (i.e. more capable of modeling the real behaviors of the range) than others, an attribute that is difficult to define. As a proxy for these two problems, we performed noisy analyses using 100 of the previously-simulated datasets by adding noise to the ellipses observed at the present. For each simulated dataset, we generated this noise based on the range of observed values at the tips for a particular character (*range* = *maximum* − *minimum*). The extra noise for each character at each tip was drawn from a uniform distribution between −0.1 *× range* and 0.1 *× range*, i.e. up to 10% of the total range of values observed in the dataset. This meant that some noisy ellipses differed substantially from the true ellipses. We then analyzed these noisy datasets and compared our estimated root states and parameters with their true simulating values.

We also investigated EMPIRE‘s sensitivity to missing species. EMPIRE only models the phylogenetic history leading to observed species in the reconstructed tree, ignoring the effect of “hidden” speciation events that are caused by extinction or imperfect taxon sampling. Using 100 of the previously-simulated datasets, we removed 50% of observed species to represent a scenario where 50% of species either go extinct before the present, or are not sampled for other reasons. We then analyzed these datasets with missing species and compared our estimated root states and parameters with their true simulating values.

Of all current biogeographic models, EMPIRE is most similar to *rase* (Quintero et al. 2015). The *rase* model describes the evolution of range centroids in space using Brownian motion while accounting for species’ entire ranges at the present. The advantage of *rase* is that it uses true species ranges, and can incorporate information about occurrence probabilities or density of occupancy. EMPIRE differs from *rase* because it uses ellipses to summarize ranges, but it also models these ellipses going back in time, including asymmetric cladogenetic range inheritance events at speciation. To highlight the differences between the models, we applied *rase* to datasets simulated under the EMPIRE model.

### Australian Sphenomorphinae

After validating EMPIRE, we applied it to the Australian Sphenomorphinae, a clade of approximately 280 described skink species. We used a dated phylogeny subset from the squamate tree constructed by Title et al. (2024), containing 218 species and with a root age of ~36 Ma. Expert range maps for each species were obtained from the Global Assessment of Reptile Distributions (GARD) database (Roll et al. 2017; Caetano et al. 2022). To generate an ellipse from each expert range map, we randomly sampled 10,000 points from the interior of the range polygon(s) and drew a 95% confidence ellipse around these points, then calculated ellipse characters (*x, y, r, s, a*) from the confidence ellipse. Bear in mind that latitude/longitude coordinates were treated as a flat, square grid. This is reasonable for our analyses because Australia is somewhat near to the equator, but other studies may necessitate different coordinate systems.

We ran EMPIRE for 2,000,000 generations with 200,000 generations used for burn-in to tune proposal step sizes. Priors on root values and the long-term mean of the OU process on log area of the projecting circle (*root*_*x*_, *root*_*y*_, *root*_*r*_, *root*_*s*_, *root*_*a*_, *µ*) were normal distributions centered on the empirical means of modern-day ellipses, with standard deviations equal to the range of the observed values (*maximum* − *minimum*). Priors on volatility parameters and the rate of mean reversion for the OU process on log area (of the projecting circle) were uniform distributions from 0 to 10 (*σ*_*x*_, *σ*_*y*_, *σ*_*r*_, *σ*_*s*_, *κ*) or from 0 to 40 (*σ*_*a*_).

After running EMPIRE on the skink data, we estimated root states and model parameters using their mean posterior values. To reconstruct ancestral states, we first determined the most favorable cladogenetic scenario at each node (the most frequent combination of the 4 discrete characters **w**_*node*_ = *{d, m, c, h}*), then assigned continuous ellipse characters at each node using the mean of the posterior distribution consistent with the selected scenario at that node. We also found the posterior distribution (thinned 1:500) for true (projected) directions of cladogenesis at each node, binned into four categories: E-W splits, NE-SW splits, N-S splits, and NW-SE splits. Note that ‘splitting direction’ refers to the major axis of movement for any speciation event; a N-S split refers to any event that results in one daughter centroid being located (approximately) north of the other. Both ‘splitting’ and ‘budding’ type events have a ‘direction of cladogenesis’. We compared these directions across three major clades: the two large genera, *Ctenotus* and *Lerista*, and a third clade containing all but one remaining species. We also mapped the most common directions of cladogenesis for ellipses with centroids in different grid cells across Australia.

Next, we investigated how the aridification of Australia may have contributed to the evolution of these skinks. We used 20-year modern precipitation data from the Australian Government Bureau of Meteorology (2024), and paleoclimate data at 20 Ma and 40 Ma from Pohl et al. (2022), reoriented to the coordinate system of modern Australia using the PALEOMAP model with *RGplates* (Roll et al. 2017; Caetano et al. 2022). From these data rasters, we generated two new rasters, one representing the average change in precipitation *mm × day*^−1^ *× MY* ^−1^ during the period from 0–20 Ma, and the other representing the average change in precipitation during the period from 20–40 Ma. These rasters provide a simple view of Australia’s changing climate over an extensive time period, but comprehensive data at a finer scale is not available.

To determine whether aridification was associated with increased speciation, we constructed Poisson regression models for the 0–20 Ma time period; the older time period had too few speciation events to perform a meaningful regression. We modeled the log of the number of speciation events involving a grid cell (the number of reconstructed ancestral ellipses overlapping that grid cell) as a function of an intercept term and up to two predictors: the change in aridity in *mm × day*^−1^ *× MY* ^−1^, and the distance to the nearest continental boundary (scaled to have mean 0 and standard deviation 1). This was done using the spind R package, which uses wavelet-revised methods to account for spatial autocorrelation in both the predictor variables (aridity and edge distance) and the response variable (speciation event count) (Carl et al. 2018). This is important because we expect spatial autocorrelation in precipitation and species counts, regardless of their relationship to one another. As *spind* requires a square grid, climate rasters were resampled using a square grid system.

To determine whether daughter ellipses demonstrated ecological partitioning, we compared the difference in mean aridity between the two daughter ellipses under the reconstructed scenario at each node to the mean difference in mean aridity across all possible scenarios for the same ancestral ellipse.

## Results

### Simulations

Simulation results indicate proper coverage across all root states and model parameters. Mean coverage was approximately 95% across root states and model parameters, with all coverage values falling between 93% and 97% (Fig. 3). The median effective sample size (ESS) for all analyses across root states and model parameters was approximately 1300, where 200 is usually considered sufficient for accurate estimates. Point estimates (mean posterior values) also align well with the true simulation values. Rate parameters governing the movement, elongation, and expansion of ellipses are the easiest to estimate, especially the *µ* parameter governing the long-term mean of the OU process describing range growth dynamics. Root values for elongation characters are also well estimated, but the root values for the ancestral centroid location and size are somewhat more diffuse. These are the ellipse characters which change at cladogenesis, so this is somewhat expected; examining many potential histories of cladogenetic events creates wider HPD intervals and less precise point estimates. However, the model is correctly exploring these options, as coverage values are excellent. Also, due to the way that range size is modeled (an OU process on log area of the projecting circle) and the sizes of some new ranges immediately after speciation (*α*), it is not possible to estimate very small range sizes for the MRCA (*root*_*a*_) with sizes between *e*^−5^ ≈ .007 and *e*^0^ = 1. While these values are dependent on the scale and parameter values of our particular simulations, they suggest that very small ranges, especially those close to the value of *α* and for older cladogenetic events, may be difficult to estimate. In general, EMPIRE‘s ability to estimate root states and model parameters does not depend strongly on tree size, although larger trees do provide slightly better estimates of rate parameters (Supplemental Fig. 3).

### Australian Sphenomorphinae

Model parameters governing the movement, elongation, and cladogenetic inheritance of range ellipses were estimated using the mean of the posterior distribution for each parameter (Fig. 4). In this analysis, distances were measured in degrees latitude or longitude, and time was measured in millions of years. Note that latitude/longitude coordinates were treated as a unbounded, flat, square grid.

**Figure 4.**
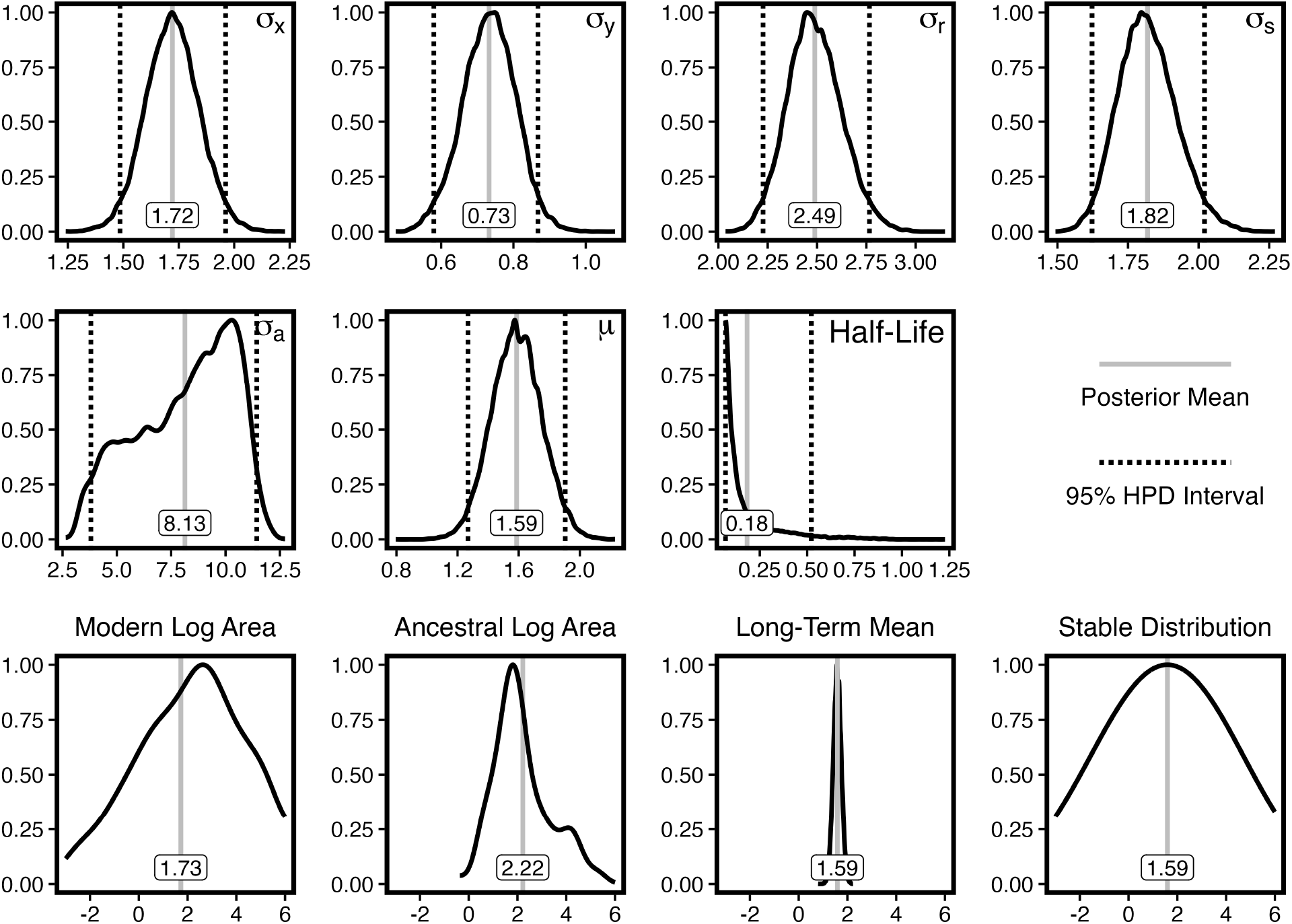
Estimates for root states, model parameters, and ellipse sizes for the Australian Sphenomorphinae dataset. Distances are measured in degrees latitude or longitude and time is measured in millions of years.

Mean posterior estimates for parameters governing the volatility (i.e. the square root of the diffusion rate) of movement of range ellipse centroids, *σ*_*x*_ and *σ*_*y*_, have values of 1.72 and 0.73 degrees/Myr, respectively (~190 kilometers and ~80 kilometers, assuming 1 degree ~111 kilometers). As *σ*_*x*_ describes east-west motion and *σ*_*y*_ describes north-south motion, we can conclude that the skinks’ range ellipses tended to move east-west ~58% more than north-south. Similarly, the volatility of east-west change of the tilt point controlling oblongness, *σ*_*r*_, has an estimated value of 2.49 degrees/Myr, while the volatility of north-south change, *σ*_*s*_, has an estimated value of 1.82 degrees/Myr. Recall that these rates apply to the tilt point that controls oblongness, and therefore do not have a direct geographic interpretation; however, their relative values suggest that expansion in the east-west direction was also less limited than expansion in the north-south direction. Although a shorter physical distance is required to traverse 1 degree longitude than 1 degree latitude due to the curvature of the Earth, this does not account for the estimated differences in rates of range movement or range elongation. Additionally, a substantial amount of movement was attributable to cladogenetic events that resulted in a relocation of the ellipse centroid; the total amount cladogenetic movement was ~39% of the total anagenetic movement. Of all reconstructed cladogenetic events, ~63% were budding events, and ~37% were splitting events.

The OU process on the area of the projecting circle indicates a long-term mean value of ~1.59 log square degrees (this corresponds to a minor radius of ~140 kilometers for range ellipses) and a log-area half-life of ~0.18 MY, with a variance of ~8.13. Because many small ranges are created after cladogenetic events, this suggests that there is a strong tendency for ellipses to grow in size, with a substantial amount of variation in their specific growth dynamics. Reconstructed ancestral ellipses were slightly larger than the ellipses for extant species, having a posterior mean log area (of the projecting circle) of ~2.22 compared to ~1.73. This is consistent with the expectation that ellipses will tend to grow in size, and will speciate when they are large.

Overall, reconstructed ellipse centroids remain predominantly within Australia, and reconstructed ellipses appear similar to modern-day range ellipses (Fig. 5, Supplementary Fig. 8). We did find one exception: an interior node within *Ctenotus* with three present-day descendants. This ellipse is abnormally large and north-south elongated. Examining the descendant nodes, we can see that this large ellipse is caused by the east-coast location of its left daughter (*Ctenotus allotropis*) and west-coast location of its right daughter (the ancestor of western species *Ctenotus australis* and *Ctenotus alleni*). This is the only reconstructed ellipse that far exceeds the bounds of the continent, and it probably only takes this shape because both daughters are distant and north-south elongated. Any other configuration (a smaller ancestor or rounder ancestor) would require large movements in (*x, y*) or (*r, s*) that are easier to account for with size. Clearly, this ancestral range and the dynamics of its descendant lineages are poorly described by this reconstruction, as skinks do not actually live in the ocean.

**Figure 5.**
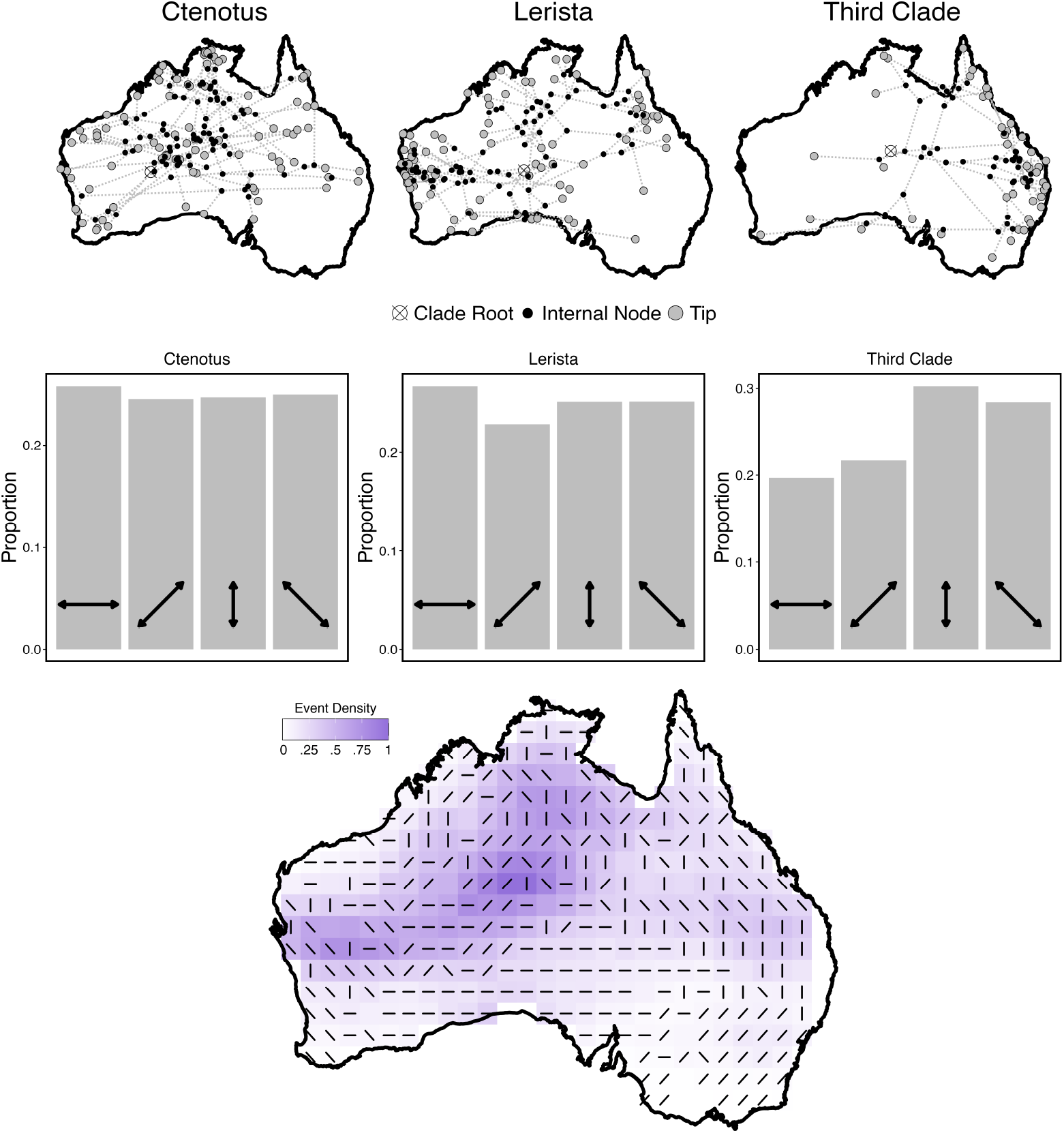
**Top:** Centroid locations of present-day species, ancestral nodes, and MRCAs for each major clade in the Australian Sphenomorphinae dataset: large genera *Ctenotus* and *Lerista*, and a third clade encompassing all but one remaining species. **Center:** Posterior distribution of binned directions of cladogenesis across all nodes by clade. **Bottom:** Most frequent binned direction of cladogenesis by grid cell. Missing cells indicate that two or more directions were equally favored. Shading indicates the number of reconstructed speciation events overlapping the grid cell.

The posterior sample of directions of cladogenesis across all nodes, binned by major clade, shows differences between the two large genera (*Ctenotus* and *Lerista*) and the third clade containing all but one remaining species (Fig. 5). The third clade, which is concentrated in the east, demonstrates far more north-south splits than east-west splits. This pattern is not observed in either *Ctenotus* or *Lerista*. This makes sense in light of the third clade’s distribution along the eastern coast of the continent. We observe that many present-day species and reconstructed ancestors are elongated along coastlines. Therefore, splits parallel to the coastline, where daughters are re-centered to disparate coastal locations, are more consistent with isolated daughter lineages. A raster showing the primary direction of cladogenesis in each grid cell across Australia (constructed using the centroid locations of each speciating ancestor across the whole posterior distribution for all nodes) demonstrates that ellipses consistently split parallel to the coastlines. When the structure of the phylogeny is ignored (i.e. the range ellipses at the tips are shuffled), this pattern is eliminated, suggesting that this is not a *post hoc* artifact of the shape of the continent, but a result of real splitting patterns among the Australian Sphenomorphinae.

Complementing the clade-wise summary of ancestral ranges, as centroids, in Figure 5 (top), we also produced more detailed probabilistic representations for the ancestral ranges, as ellipses, for each nodes in the phylogeny, individually. Figure 6 displays the ranges before and after an ancestral species gave rise to the daughter lineages that would ultimately contain *Ctenotus uber* and *C. rutilans* in one subclade, and *C. hilli, C. militaris, C. pulchellus, C. gagudju*, and *C. kurnbudj* in the other. In this case, the ranges of the ancestor (node 235) and left daughter (node 240) are large, congruent, and centered in western Australia, whereas the right daughter (node 236) is a smaller, budding range restricted to northern Australia (see Supplemental Fig. 7 to reference against the phylogeny).

**Figure 6.**
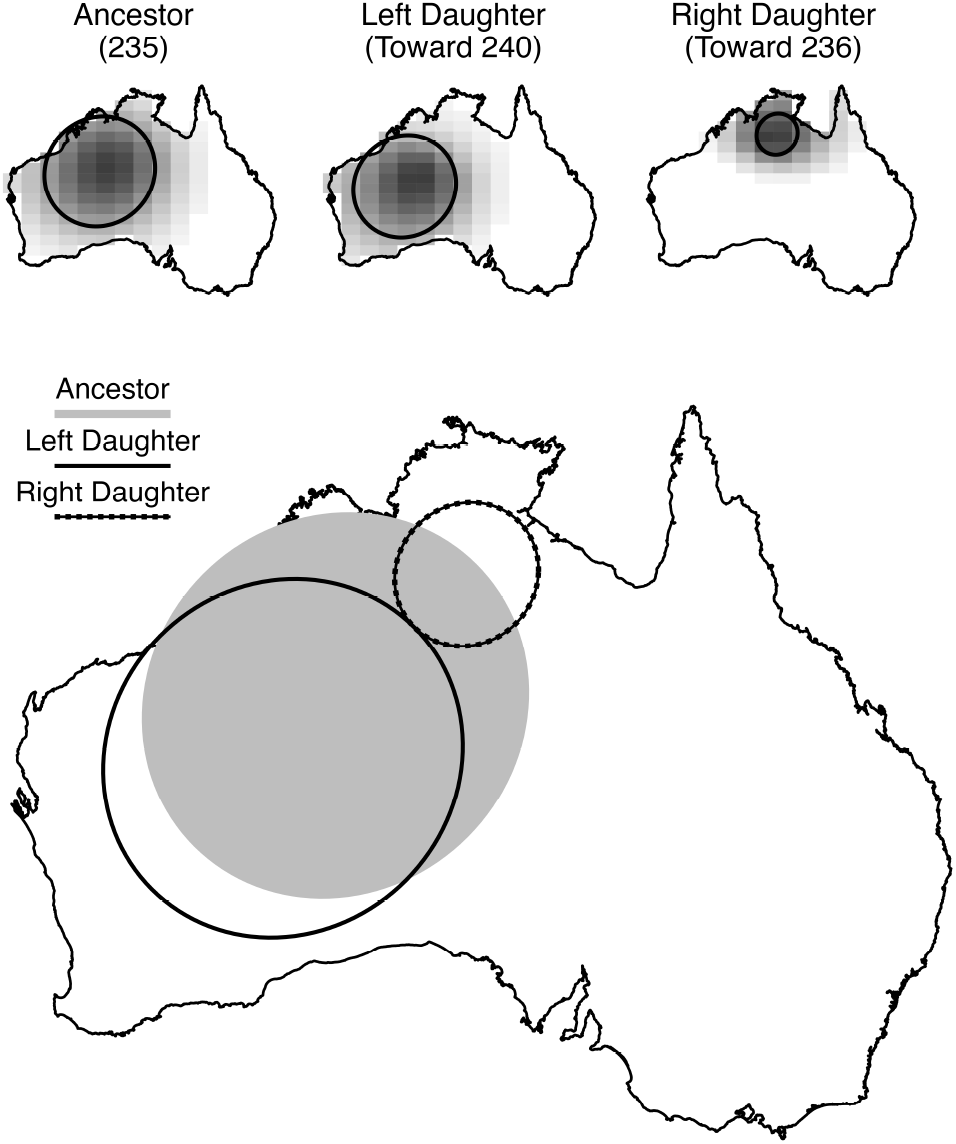
An example of a posterior distribution for a speciation event at a node (node 235, see Supplementary Figure 7 for node labels). To construct the gridded posterior distributions for ancestor, left daughter, and right daughter ellipses (top), data-augmented values for each MCMC sample were converted into ellipses. Grid cell shading represents the frequency of ellipses overlapping each cell. To reconstruct the most probable event (bottom), the most frequent combination of discrete options **w**_*node*_ = *{d, r, s, a}* was selected, and mean values for **v**_*node*_ = *{r, s, a}* were calculated, conditioned on the discrete scenario.

**Figure 7.**
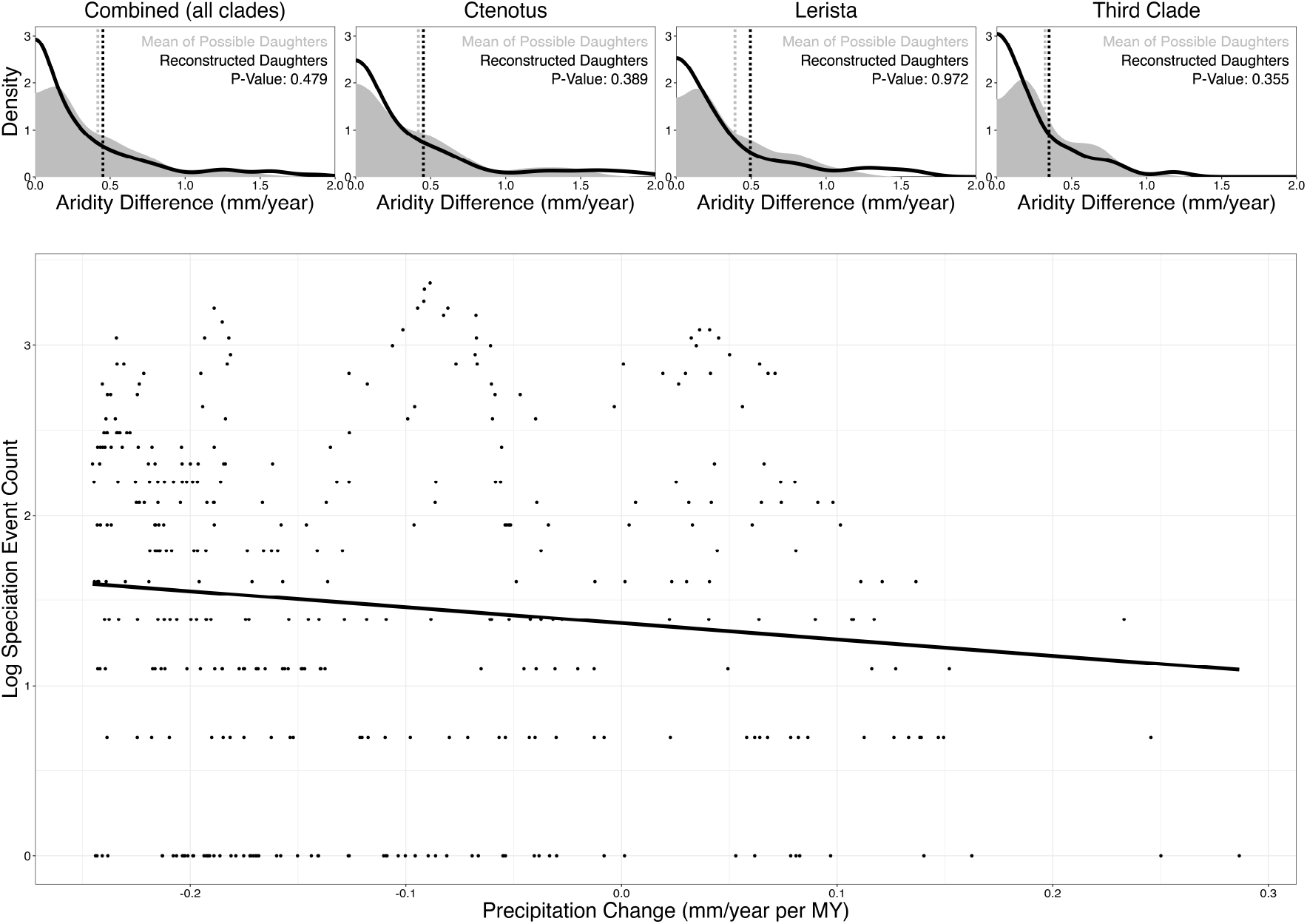
Relationships between Australian paleoclimatic dynamics and Sphenomorphinae biogeography. **Top:** Difference in mean aridity between reconstructed daughters compared to the average difference in mean aridity among all possible scenarios involving the same ancestor. Reconstructed daughters do not demonstrate greater ecological partitioning than random daughters. **Bottom:** Relationship between precipitation changes and the log of the speciation event count by grid cell (the number of reconstructed ancestral ellipses overlapping the grid cell). Here, uncorrected datapoints (without accounting for spatial autocorrelation or other predictors) are shown. Aridification is positively associated with speciation.

It has also been suggested that changes in aridity across Australia during the evolution of the Australian Sphenomorphinae may have contributed to their diversification. To examine whether reconstructed ancestor and daughter lineages demonstrated ecological partitioning, we assessed whether reconstructed daughters were more different from one another than randomly sampled daughters. We did this by comparing the distribution of differences in mean aridity among reconstructed daughters to the distribution of average differences in mean aridity among all possible daughters arising from the same reconstructed ancestors. For all Australian Sphenomorphinae and each individual clade, we did not find evidence of ecological partitioning (Fig. 7). This pattern held when we examined all reconstructed ellipses, sister pairs alone, and centroids alone. It may be that there is no relationship between aridity and direction of cladogenesis, or that there is not sufficient statistical power to detect such a pattern, given the number of nodes in the phylogeny of the Australian Sphenomorphinae.

However, we did identify a relationship between aridification and speciation. Using the reconstructed ancestral ellipses, we counted the number of speciation events involving (overlapping) each grid cell across Australia, and modeled the log of the speciation event count as a function of precipitation change and distance from the nearest continental edge using wavelet-revised methods accounting for spatial autocorrelation. We found that a model incorporating both edge distance and precipitation change was preferred over a distance-only model by 21 AIC points, indicating that aridification is an important predictor of speciation in this system. A negative coefficient was estimated, indicating that more rapidly aridifying regions experienced more speciation events. For example, the grid cells showing the most aridification (*<* −0.2 *mm × day*^−1^ *× MY* ^−1^) experienced an average of 7.4 speciation events over 20 million years, while the grid cells showing the greatest precipitation increase (*>* 0.2 *mm × day*^−1^ *× MY* ^−1^) experienced an average of 2 speciation event over the same time period. A positive relationship between aridification and speciation suggests that, while daughters do not partition the changing ecological conditions, those conditions nevertheless contribute to speciation through other mechanisms. This pattern was only observed by reconstructing entire ancestral ellipses; when using centroids alone, a distance-only model was preferred.

## Discussion

Our analysis of the Australian Sphenomorphinae demonstrates the value of EMPIRE for investigating the ways in which species ranges move, elongate, grow, and divide. EMPIRE models the position, extent, and orientation of species ranges as ellipses evolving over time and moving through space, including asymmetric range inheritance events at speciation. This allows us to reconstruct ancestral range ellipses, examine the movement and growth dynamics of lineages, and assess how the spatial context of speciation is related to the environment.

Our results suggest that the continental boundaries of Australia played a major role in rates of movement and elongation. They were also the primary contributor to the direction of cladogenesis during speciation events, resulting in location-based differences in predominant directions and per-clade differences in the most common directions. Importantly, EMPIRE was able to recover these coastal influences despite the fact that boundaries are not incorporated into the model itself. Under the EMPIRE model, space is assumed to be flat, never-ending, and traversable. Therefore, the estimated differences in rates of movement and elongation in each cardinal direction, the reconstructed locations of ellipses, and the alignment of the directions of cladogenetic events to the coastlines are the exclusive result of the provided phylogeny and spatial data about present-day species ranges. Even if we did not know the boundaries restricting the ranges of Australian Sphenomorphinae, we could roughly estimate them from our reconstructions. We avoid speculating about the precisely which factors cause this subtle pattern, as any number of biotic or abiotic factors, alone or in combination, could have elevated how often species originate in alignment with the continental perimeter. For example, environmental gradients often run parallel to coastlines, whereas river (barriers) are orthogonal, either of which might shape cladogenetic patterns. The fact that skinks cannot live in the ocean is also relevant.

As expected, range size dynamics showed a strong tendency toward growth. Under some of EMPIRE‘s cladogenetic scenarios, one or both daughter ranges may be greatly reduced after speciation. The OU process on the log area of the projecting circle allows a reduced range to reach a larger size prior to the next speciation event. Reconstructed ancestral range ellipses were somewhat larger than present-day range ellipses, indicating that large ranges are associated with speciation.

We were also able to investigate the relationship between the changing environment of Australia and the diversification of the Australian Sphenomorphinae. We found that speciation events were positively correlated with aridification when accounting for the spatial autocorrelation of both speciation events and precipitation conditions. This pattern only emerged when using whole reconstructed range ellipses; centroids alone were not sufficient to demonstrate this pattern. By reconstructing the daughter ellipses immediately after speciation, we found that the daughters did not partition the new environmental conditions. Daughter ellipses were not more different than the average of all possible daughters arising from the same ancestor. Instead, alternative explanations are implicated, such as reduced population sizes, range fragmentation, or isolation resulting from range expansion.

Here, it is critical to reiterate that range ellipses are summaries of the extent, position, and orientation of species ranges; they do not represent range boundaries, and it should not be assumed that a species will occupy the entirety of its range ellipse. This means that reconstructed speciation events may represent a variety of allopatric, peripatric, or sympatric scenarios depending on the actual distribution of individuals within the range ellipse. Consider, for instance, a species with a disjoint range comprised of several spatially-disparate populations. Under the budding speciation mode, EMPIRE explores the possibility that a new daughter lineage may arise entirely within the range ellipse of the ancestor. While this may initially seem like sympatry, it could equally describe a situation where one of the isolated populations becomes differentiated, an allopatric event. We caution against using the results of an EMPIRE analysis to draw conclusions about the connectivity of populations within ellipses.

Similarly, the two speciation event types presented here do not encompass all possible speciation events that one might imagine. Speciation along a gradient, for example, might result in two overlapping daughter species of similar size. It would be relatively straightforward to incorporate additional modes into the EMPIRE model, or additional grid options for concentric circles and direction lines. However, we argue that adding more detail to speciation events may result in diminishing returns. EMPIRE does not recover discrete scenarios with a high degree of confidence. Even if each speciation event left a strong spatial signal on the distribution of modern species (which is not the case), we would still expect EMPIRE to assign high probability to several similar scenarios. EMPIRE effectively explores discrete options at each node, generating relatively smooth posterior distributions of ancestor and daughter lineages. These posterior distributions most effectively display the spatial context of a speciation event, as well as the uncertainty associated with the event. Some events demonstrate dramatic spatial differences between daughter lineages, while others do not.

In our aridity analyses, we used reconstructed speciation events based on the most frequent scenario (combination of discrete options) and the mean values of continuous characters under that scenario. These reconstructions appear to represent the posterior distributions of ancestors and daughters well, and are simpler to obtain and summarize. However, the EMPIRE package stores all MCMC output, including data-augmented values at nodes, in simple file formats. This allows users to examine the complex posterior distribution in whichever way they prefer.

Effectively exploring the possible combinations of values for parameters and data-augmented characters at nodes can be a challenge. The EMPIRE package uses a set of relatively simple MCMC moves to accomplish this. By default, some move types are scheduled more frequently than others (weighting). These weights were chosen to improve mixing for trees of sizes between 20 and 250 tips. For larger phylogenies, it will be helpful to increase the weights for proposals on data-augmented values, i.e. ellipse characters at nodes. Additionally, if a burn-in period is assigned, these iterations are used to auto-tune the proposal widths for each move type. This is necessary for proper mixing. Keep in mind that these iterations are saved by default, and should be removed during output processing, as they do not correctly represent the posterior.

As expected, sensitivity analyses representing scenarios with noise added to ranges for extant species (Supplemental Fig. 2) indicate that substantial amounts of noise (up to 10% of the full range of values in the dataset) typically cause volatility to be overestimated. This is because additional variation in ellipse characters is being introduced at the tips of the phylogeny. However, the deterministic part of the OU process on the log area of the projecting circle, particularly the long-term mean of the process *µ*, is still estimated accurately.

Large numbers of missing taxa (50% of simulated taxa removed) also cause the overestimation of volatility (Supplemental Fig. 1) because character change, which would normally be attributed to the missing cladogenetic events, must instead be attributed to the Brownian and OU motion of ellipse characters. Other continuous-space biogeographic models that do not incorporate centroid relocation at cladogenesis show similar behavior. For example, when we applied the *rase* model (which does not incorporate asymmetrical cladogenetic events) to data generated by EMPIRE, rates of centroid movement were overestimated (Supplemental Fig. 4). This was especially true when ellipses were large (Supplemental Fig. 5). We note that these sensitivity analyses were designed to test the performance of EMPIRE under analysis conditions that are substantially different from the proposed data-generating process. In real, empirical systems, we expect the degree of misspecification to be less severe. We offer these findings primarily as a caution for those studies where the amount of measurement error or the number of unsampled or extinct taxa is known to be large.

In the future, extensions of the EMPIRE model may address some of its current limitations, especially those involving the assumption of a single, known, dated phylogeny without missing taxa. The first step may be incorporating additional taxa with unknown phylogenetic positions or ranges using interpolation and averaging, and potentially performing data augmentation along the branches of the phylogeny to account for hidden speciation events. The addition of a molecular model for joint divergence time dating would also be feasible; however, uncertainty in phylogenetic relationships would be more difficult to model. If a posterior sample of trees is already available, one solution would be to perform multiple EMPIRE analyses on different trees from the posterior sample and compare the results.

It also may be possible to incorporate environmental features and barriers into the model itself. The current implementation of EMPIRE uses simple Brownian motion to describe the movement and elongation of species ranges. However, bounded or asymmetric processes could be examined. Alternatively, machine learning approaches could bypass the need for a likelihood function, relying exclusively on simulation.

Finally, we emphasize that modeling the evolution of ellipses and their behaviors at cladogenesis has applications to other problems that interest biologists. Ellipses can be used to describe many two-dimensional species attributes, including pairs of niche variables or correlated morphological characters. The details of anagenetic evolution and cladogenetic events may differ between problems, but EMPIRE‘s dimension-reduction and data-augmentation techniques would remain effective for reconstructing ancestral states and estimating evolutionary rates. Similar techniques may even be applied to higher-dimensional problems in the future.

## Acknowledgments

We are grateful to members of the Landis lab at Washington University in Saint Louis and the Computational Phylogenetics group at the University of Lausanne for feedback. We are also indebted to José Figueroa-López, Jonathan Myers, Dave Queller, and Jonathan Losos for helping us refine the model and study design.

## Supplementary Material

Data available from the Dryad Digital Repository:

http://dx.doi.org/10.5061/dryad.[NNNN].

## Data Availability

Data, supplementary material, and code associated with this paper are available via Dryad: http://dx.doi.org/10.5061/dryad.[NNNN]. The EMPIRE R package is also available on GitHub: https://github.com/sswiston/empire.

## Supplemental Figures

**Supplementary Table 1.** Model characters and parameters. Here we provide a list of characters and parameters related to range ellipses, their anagenetic evolution, and their behavior at cladogenesis. We also provide the projection equations used to determine the new ellipse character values following different speciation events.

| Ellipse Characters and Parameters |  |
| --- | --- |
| Centroid Location |  |
| $x$ | The $x$ coordinate of the center point of an ellipse. Evolves under univariate Brownian motion with volatility $\sigma_x$. |
| $y$ | The $y$ coordinate of the center point of an ellipse. Evolves under univariate Brownian motion with volatility $\sigma_y$. |
| Elongation |  |
| $r$ | The $x$ coordinate of an ellipse's tilt vector (controls the tilt plane, which determines the direction of elongation). Evolves under univariate Brownian motion with volatility $\sigma_r$. |
| $s$ | The $y$ coordinate of an ellipse's tilt vector (controls the tilt plane, which determines the direction of elongation). Evolves under univariate Brownian motion with volatility $\sigma_s$. |
| $z$ | The height ($z$ coordinate) of an ellipse's tilt vector (controls the tilt plane, which determines the direction of elongation). This value remains constant throughout an analysis. Changing this height may impact the relationship between $r, s$ motion and ellipse elongation. Values that are too small may cause problems with very elongated ellipses. The same $z$ value should be used when generating ellipses and conducting analyses. In our analyses, a value of 10 is used. |
| Size |  |
| $a$ | The log area of the circle which projects an ellipse. Evolves under univariate Ornstein-Uhlenbeck process with volatility $\sigma_a$, long-term mean $\mu$, and rate of mean reversion $\kappa$. |
| Stochastic Processes |  |
| $\sigma_x$ | The square root of the rate of evolution of the $x$ coordinate of ellipse centroids. Drawn from a prior distribution and updated with MCMC. |
| $\sigma_y$ | The square root of the rate of evolution of the $y$ coordinate of ellipse centroids. Drawn from a prior distribution and updated with MCMC. |
| $\sigma_r$ | The square root of the rate of evolution of the $r$ coordinate of ellipse centroids. Drawn from a prior distribution and updated with MCMC. |
| $\sigma_s$ | The square root of the rate of evolution of the $s$ coordinate of ellipse centroids. Drawn from a prior distribution and updated with MCMC. |
| $\sigma_a$ | The square root of the rate of evolution of the $a$ coordinate of ellipse centroids. Drawn from a prior distribution and updated with MCMC. |
| $\mu$ | The long-term mean of ellipse sizes $a$. Drawn from a prior distribution and updated with MCMC. |
| $\kappa$ | The rate of mean reversion for ellipse sizes $a$. Drawn from a prior distribution and updated with MCMC. |
| Cladogenetic Parameters |  |
| Cladogenetic Scenarios $\mathbf{W}$ | |
| $d$ | Left/right daughter assignment at a node. A value of 0 makes the left daughter $D1$ and a value of 1 makes the right daughter $D1$. |
| $m$ | The cladogenetic mode assigned to a node. A value of 0 indicates budding cladogenesis and a value of 1 indicates splitting cladogenesis. |
| $c, c^*$ | The concentric circle assigned to a node. Each concentric circle is given an index $c$, starting with 0. Each index is further associated with a value $c^*$, indicating a distance from the center of the circle relative to its radius. In our analyses, we use 4 concentric circles with indices $\{0, 1, 2, 3\}$ and values $\{0.0, 0.5, 1.0, 1.5\}$. |
| $h, h^*$ | The direction line assigned to a node. Each direction line is given an index $h$, starting with 0. Each index is further associated with a value $h^*$, indicating an angle in radians. In our analyses we use 8 direction lines with indices $\{0, 1, 2, 3, 4, 5, 6, 7\}$ and values $\{0, \frac{\pi}{4}, \frac{\pi}{2}, \frac{3\pi}{4}, \pi, \frac{5\pi}{4}, \frac{3\pi}{2}, \frac{7\pi}{4}\}$. |
| Prior Event Probabilities |  |
| $\rho_m$ | The prior probabilities of each cladogenetic mode (budding or splitting). Under the simple model used in our analyses, all events are considered equally probable; in a complex model, the simplexes of event probabilities could be updated via MCMC. |
| $\rho_c$ | The prior probabilities of each concentric circle. Under the simple model used in our analyses, all events are considered equally probable; in a complex model, the simplexes of event probabilities could be updated via MCMC. |
| $\rho_h$ | The prior probabilities of each direction line. Under the simple model used in our analyses, all events are considered equally probable; in a complex model, the simplexes of event probabilities could be updated via MCMC. |
| Continuous Values $\mathbf{V}$ | |
| $r$ | The data-augmented $r$ value of an ellipse assigned to a node immediately prior to speciation. |
| $s$ | The data-augmented $s$ value of an ellipse assigned to a node immediately prior to speciation. |
| $a$ | The data-augmented $a$ value of an ellipse assigned to a node immediately prior to speciation. |
| Starting States $\mathbf{root}$ | |
| $root_x$ | The $x$ value for the ancestral ellipse at the root of the clade. Drawn from a prior distribution and updated with MCMC. |
| $root_y$ | The $y$ value for the ancestral ellipse at the root of the clade. Drawn from a prior distribution and updated with MCMC. |
| $root_r$ | The $r$ value for the ancestral ellipse at the root of the clade. Drawn from a prior distribution and updated with MCMC. |
| $root_s$ | The $s$ value for the ancestral ellipse at the root of the clade. Drawn from a prior distribution and updated with MCMC. |
| $root_a$ | The $a$ value for the ancestral ellipse at the root of the clade. Drawn from a prior distribution and updated with MCMC. |
| Other |  |
| $\alpha$ | The $a$ value for new $D2$ ranges created under the budding cladogenetic scenario. This should be a small value, and might be negative. Under a simple model, $\alpha$ is fixed to a small value; in a complex model, it could be assigned a prior distribution and estimated using MCMC. In our analyses, a fixed value of -5 is used. |

**Supplementary Table 2.** Projection equations.

| Mode | Character | Ancestor | D1 | D2 |
| --- | --- | --- | --- | --- |
| Budding | $x$ | $x$ | $x$ | $x + f_1(r, s, a, c^*, h^*)$ |
| Budding | $y$ | $y$ | $y$ | $y + f_2(r, s, a, c^*, h^*)$ |
| Budding | $a$ | $a$ | $a$ | $\alpha$ |
| Splitting | $x$ | $x$ | $x + f_5(r, s, a, c^*, h^*)$ | $x + f_3(r, s, a, c^*, h^*)$ |
| Splitting | $y$ | $y$ | $y + f_6(r, s, a, c^*, h^*)$ | $y + f_4(r, s, a, c^*, h^*)$ |
| Splitting | $a$ | $a$ | $a + f_7(c^*)$ | $a + f_8(c^*)$ |

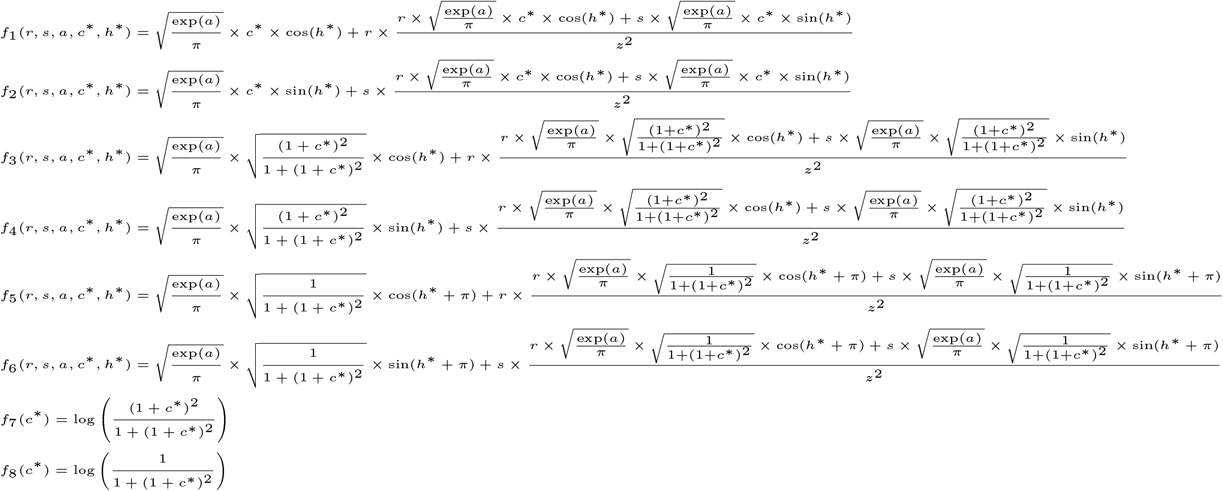

**Supplementary Figure 1.**
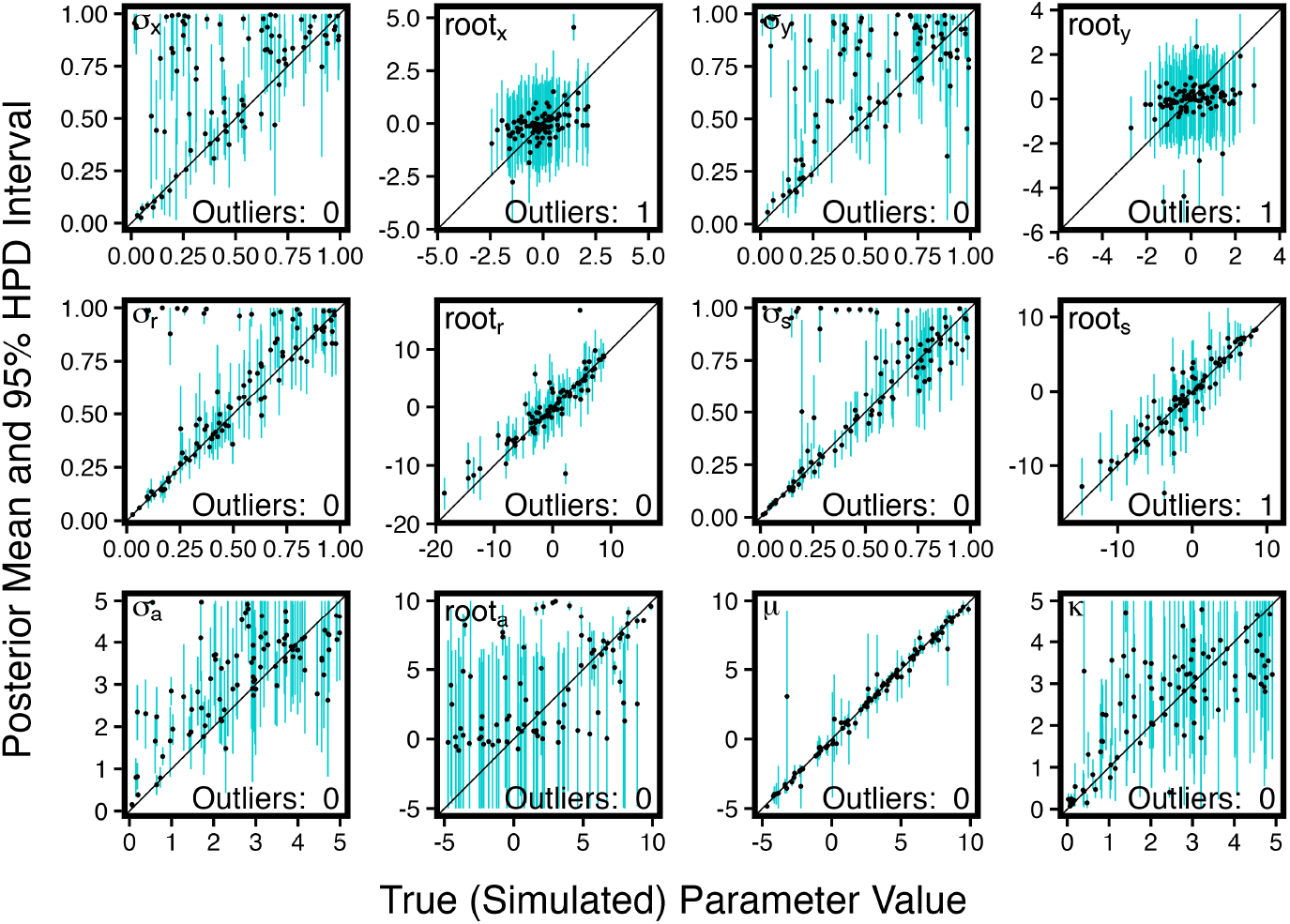
Sensitivity to missing taxa.

**Supplementary Figure 2.**
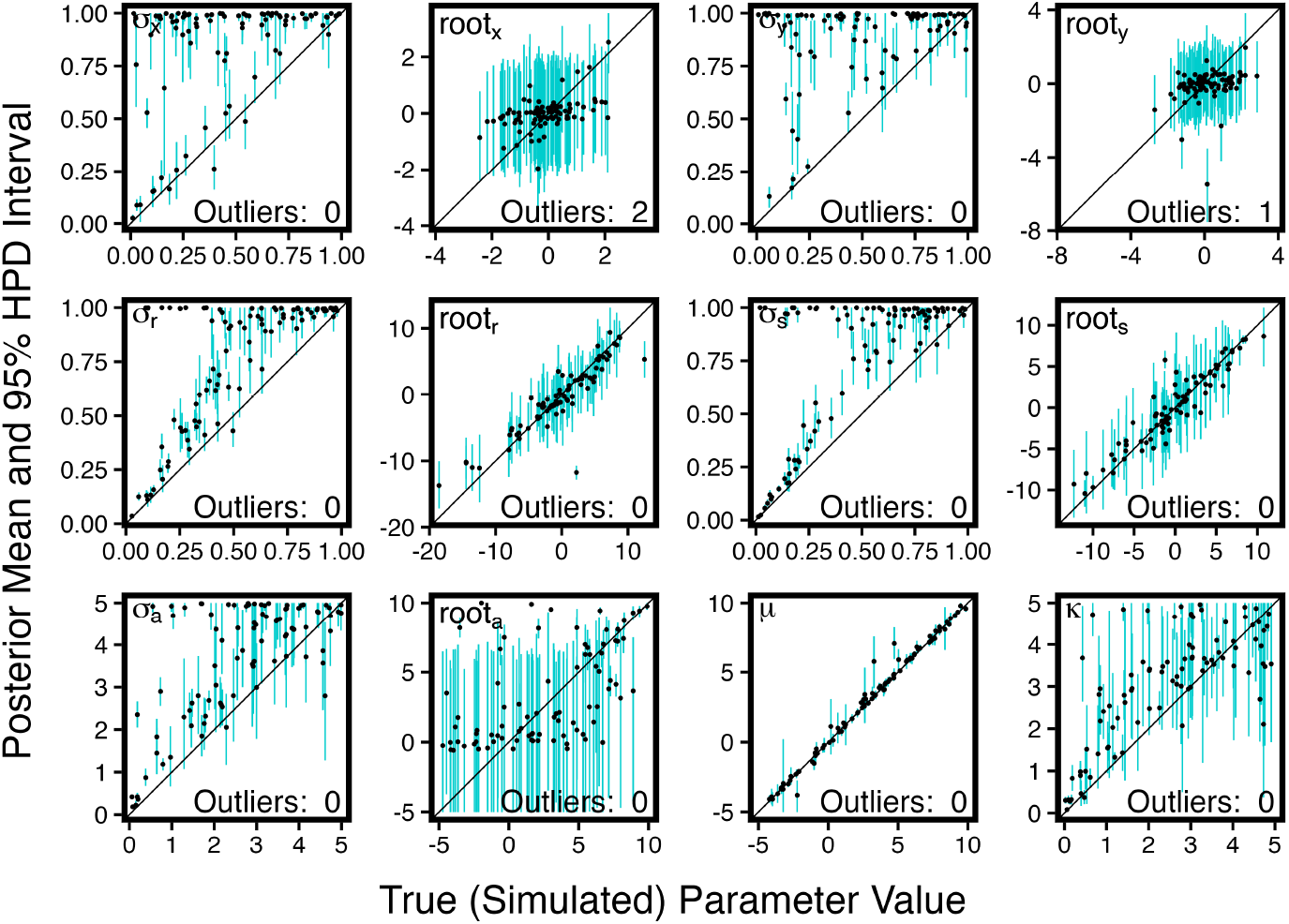
Sensitivity to ellipse noise.

**Supplementary Figure 3.**
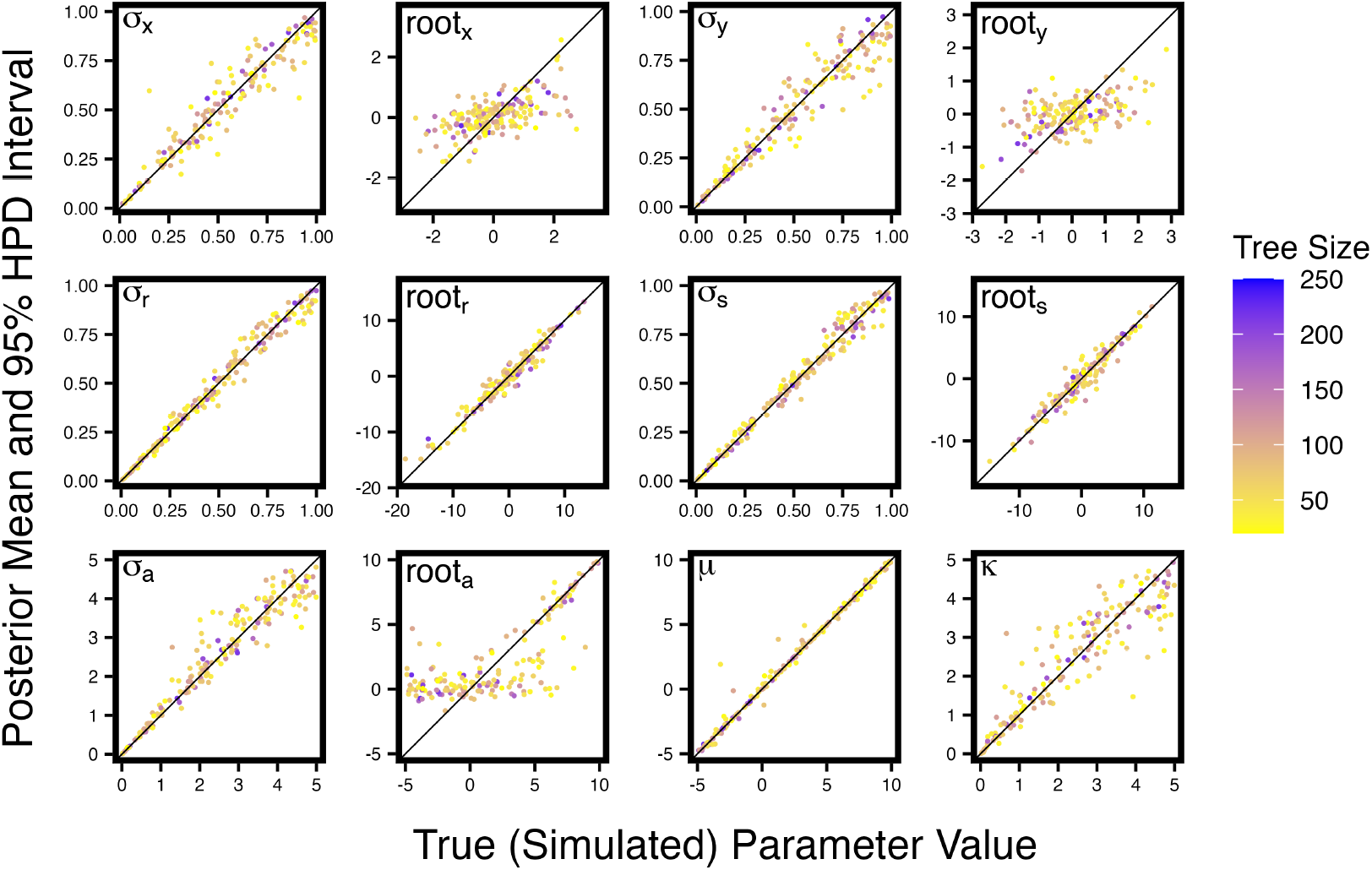
Accuracy according to tree size.

**Supplementary Figure 4.**
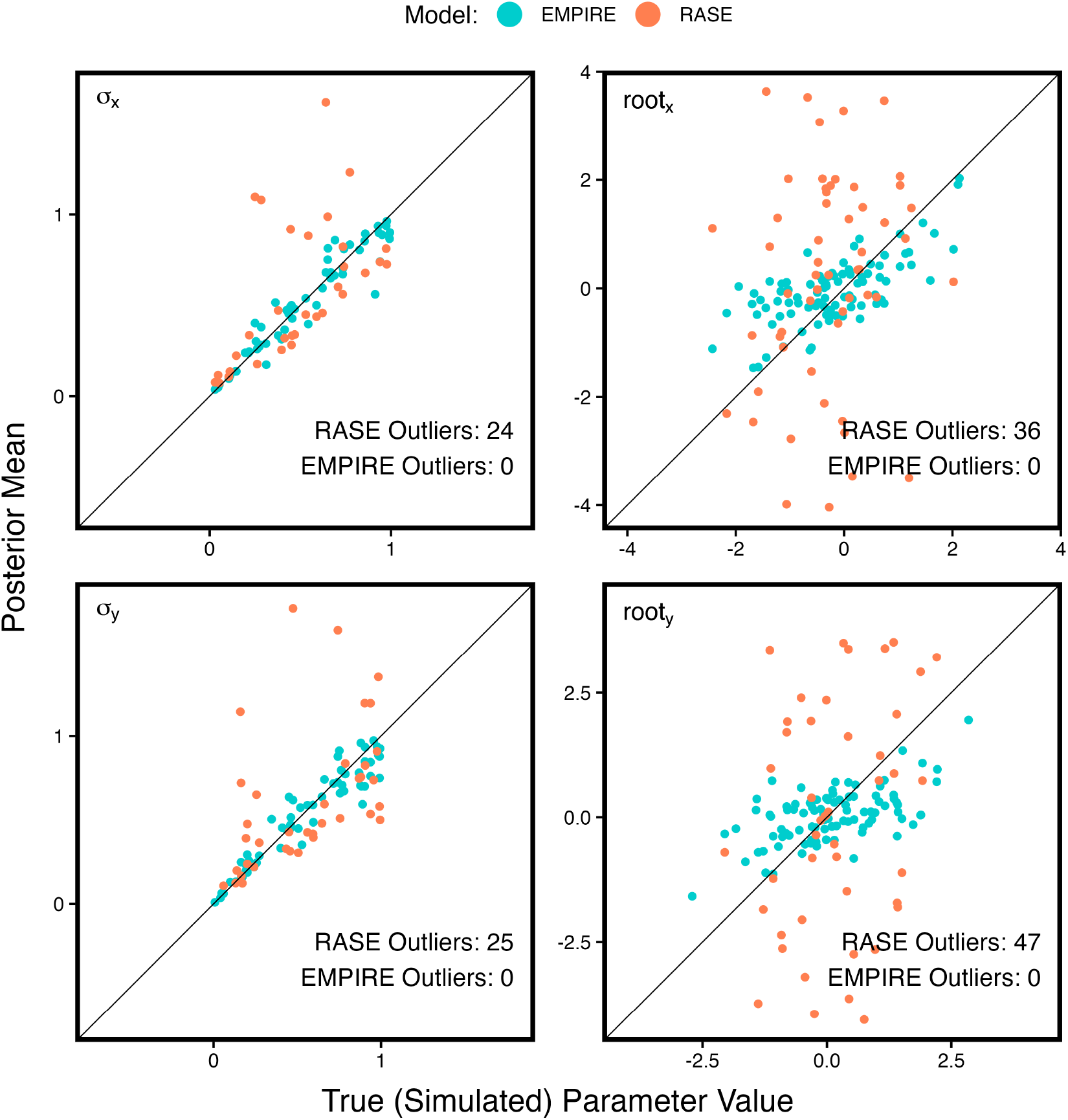
Estimates by *rase* and EMPIRE on data simulated by EMPIRE.

**Supplementary Figure 5.**
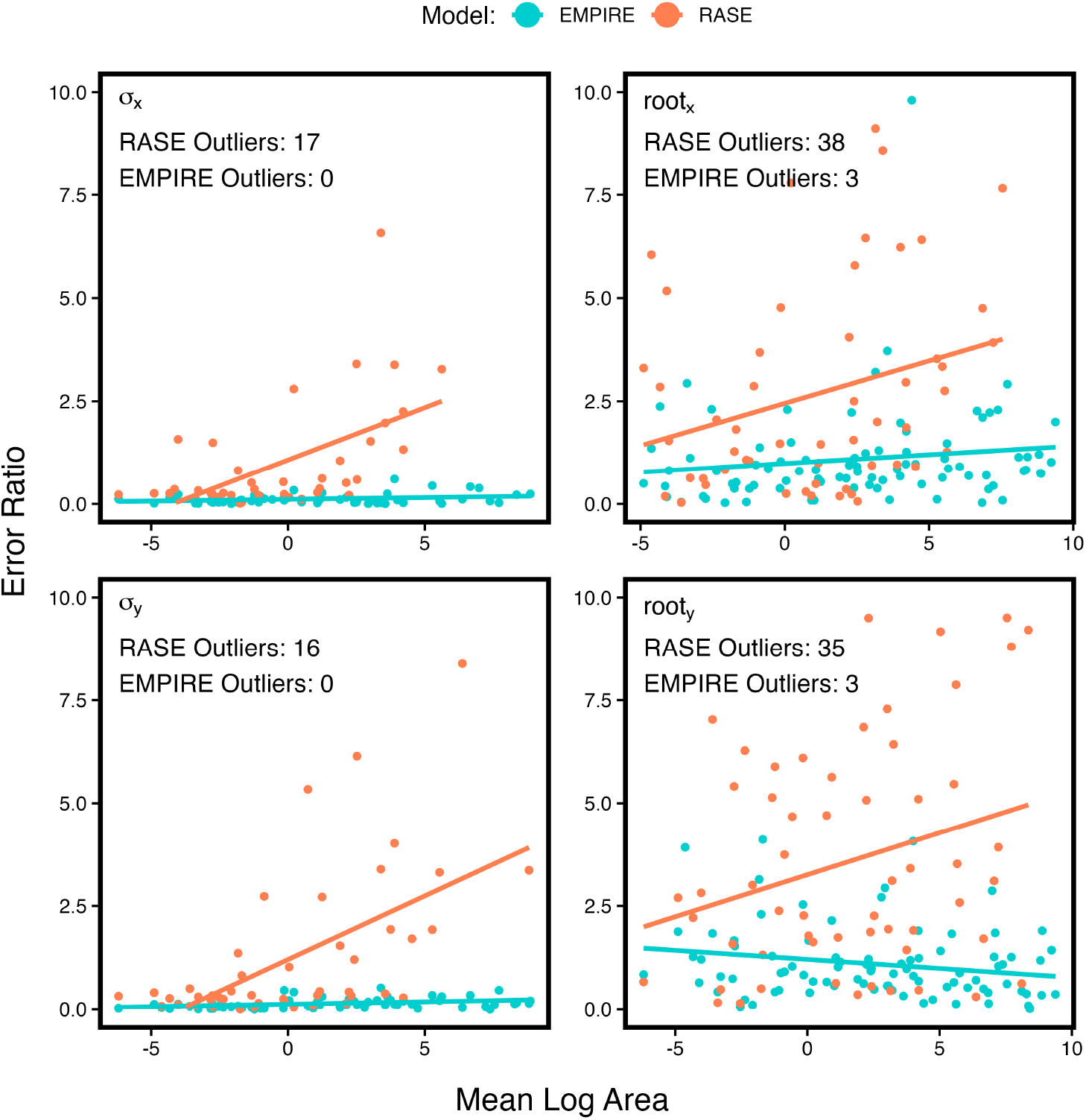
Ratio of error to true value for *rase* estimates and EMPIRE estimates on data simulated by EMPIRE, relative to the mean log area of projecting circles for reconstructed ellipses.

**Supplementary Figure 6.**
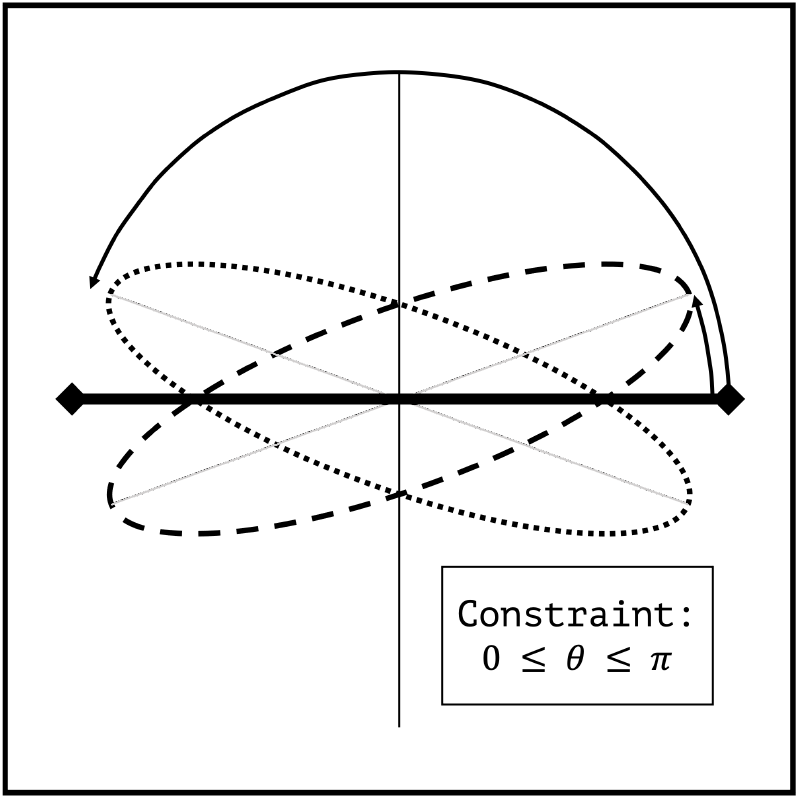
The problem with constraining rotation.

**Supplementary Figure 7.**
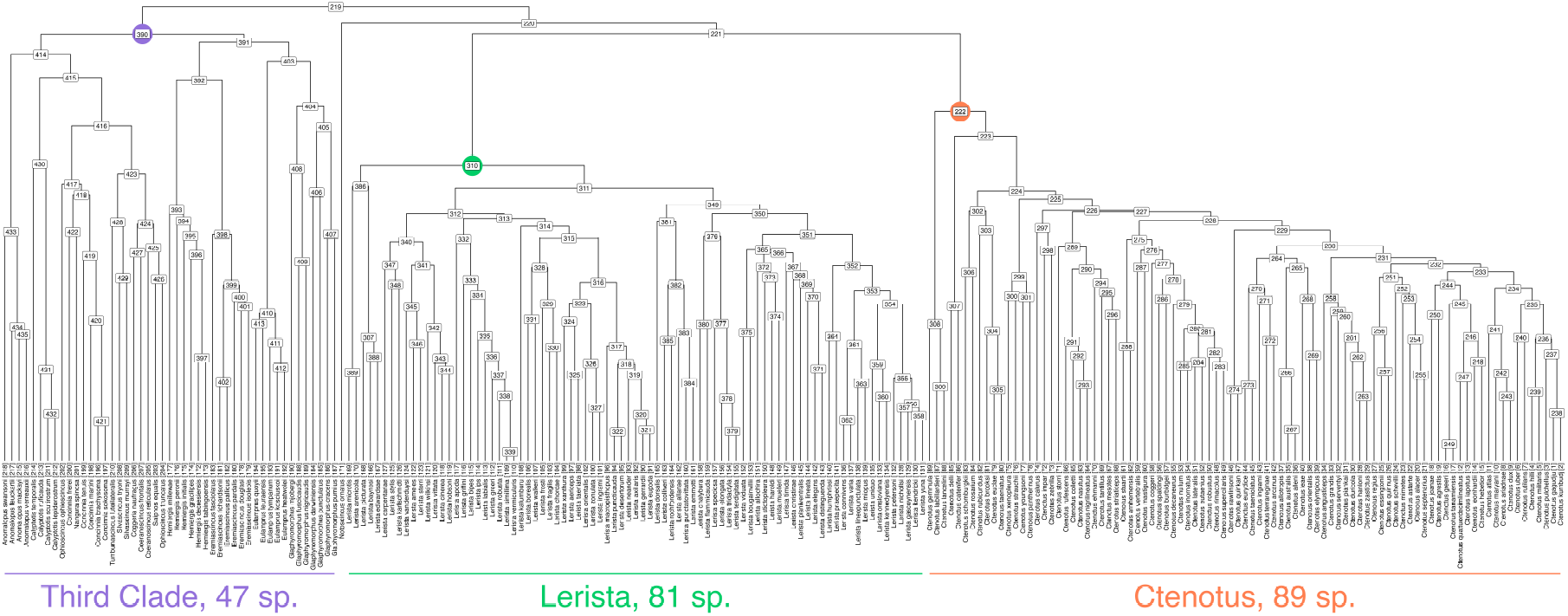
The phylogeny used for our analysis of Australian Sphenomorphinae, including node labels for internal nodes and tips.

**Supplementary Figure 8.**
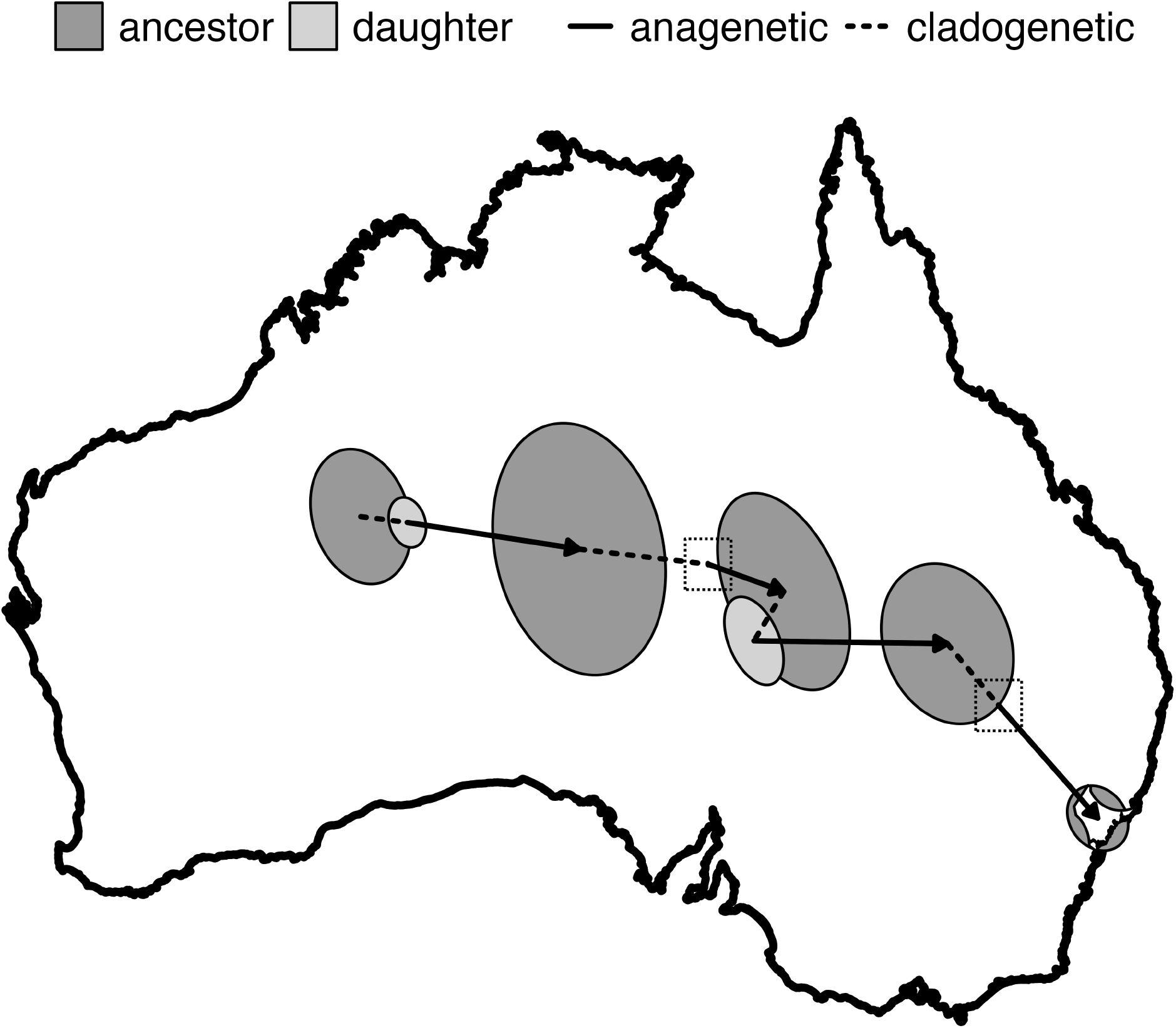
The history of a single lineage, from the common ancestor of the Australian Sphenomorphinae to the present-day species *Anomalopus swansoni*. Boxes are drawn around very small ranges for clarity.

